# PAQR5 Membrane Progesterone Receptor Regulates the Blood–Brain Barrier during Brain Metastasis Formation

**DOI:** 10.64898/2026.01.26.701753

**Authors:** Csilla Fazakas, Attila G. Végh, Tamás Dudás, Dorina Varga, Adél Lüvi, Mónika Krecsmarik, András Dér, Attila E. Farkas, István A. Krizbai, Imola Wilhelm

## Abstract

Brain metastases are a common and often fatal complication of certain cancer types, such as triple-negative breast cancer. However, the molecular pathways driving brain metastasis formation, including the migration of cancer cells from the bloodstream to the brain parenchyma across the blood–brain barrier, are not yet fully defined. Therefore, using highly relevant mouse and human model systems, the mechanisms by which triple-negative breast cancer cells and their released extracellular vesicles modulate the blood–brain barrier-forming endothelium to increase its permissiveness to tumour cell entry into the brain are investigated. It is observed that extracellular vesicles derived from tumour cells are taken up by cerebral endothelial cells, where they induce miR-146a-5p- and TGF-β1-mediated downregulation of PAQR5/mPRγ, a membrane progesterone receptor. This, in turn, leads to disruption of interendothelial tight junctions, particularly through repression of claudin-5 expression, a critical protein for maintaining barrier function. Altogether this identifies a novel mechanism by which triple-negative breast cancer-derived extracellular vesicles compromise blood–brain barrier integrity, thereby facilitating transendothelial migration of cancer cells and promoting brain metastasis development. Moreover, this study is the first to highlight the role of membrane progesterone receptors in regulating the blood–brain barrier.

**Table of contents:** Extracellular vesicles from triple-negative breast cancer cells induce miR-146a-5p- and TGF-β1-mediated downregulation of PAQR5/mPRγ, a membrane progesterone receptor, in blood–brain barrier-forming endothelial cells. This results in disruption of interendothelial tight junctions, thereby promoting enhanced migration of cancer cells into the brain. This mechanism highlights the role of membrane progesterone receptors in regulating the blood–brain barrier.

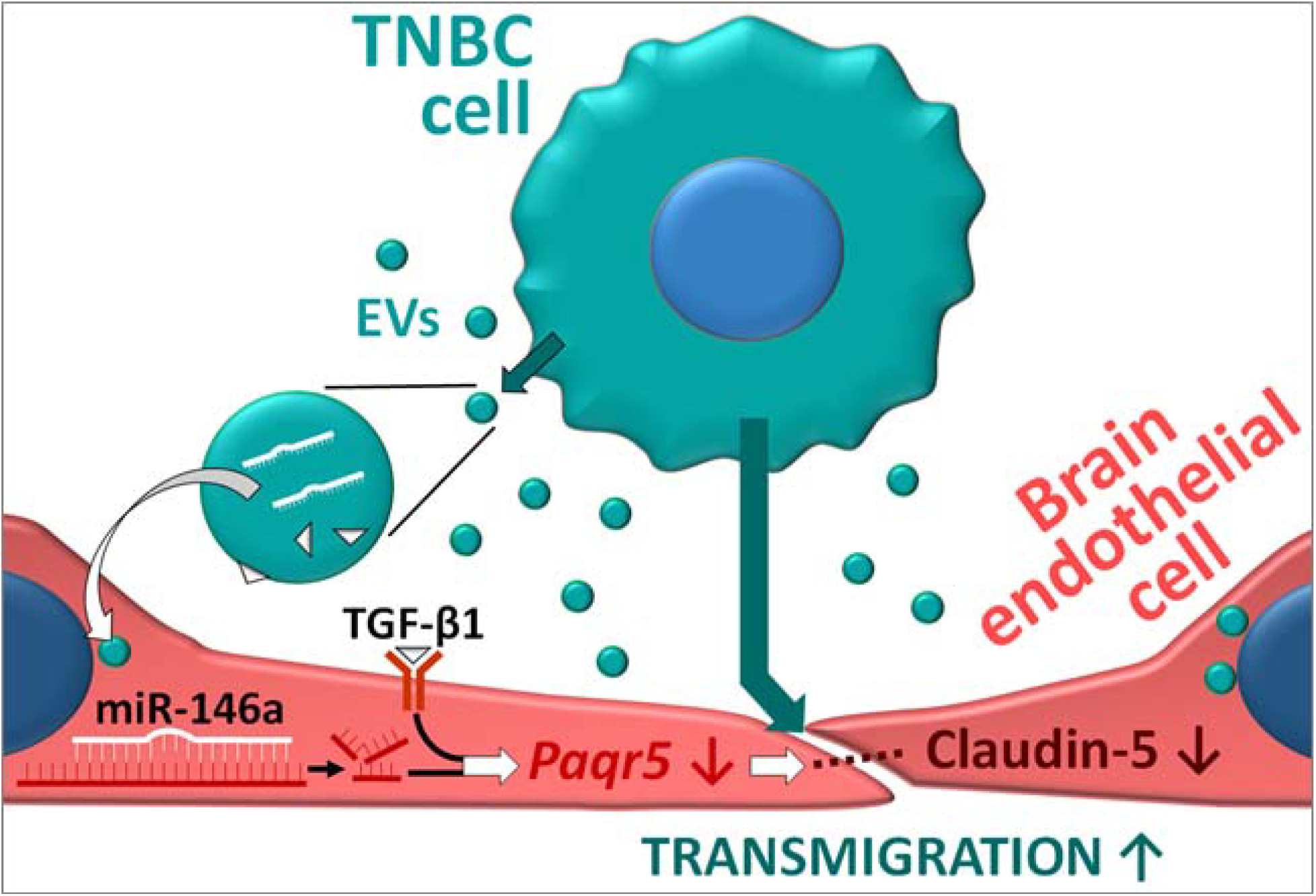

## Introduction

The mechanisms of brain metastasis formation, especially the migration of cancer cells through the blood–brain barrier (BBB), are not yet fully characterised. This lack of knowledge contributes significantly to the absence of specific therapies that could improve the dismal prognosis of the disease, as reflected in the median overall survival of less than 6 months after a brain metastasis diagnosis (Yri et al., 2025).

One of the main sources of secondary brain tumours is triple-negative breast cancer (TNBC), a molecular subtype that lacks expression of the oestrogen receptor, progesterone receptor, and human epidermal growth factor receptor type 2 (Her2) (also called Neu or ErbB-2 in mice). These tumours are usually high-grade, invasive basal-like carcinomas showing rapid growth and are more likely than other types of breast cancer to metastasise to the central nervous system (CNS) (Foulkes et al., 2010). To reach the brain parenchyma, metastatic TNBC cells have to cross the walls of brain capillaries, primarily the BBB-forming cerebral endothelium, the main role of which is to hinder the penetration of potentially toxic substances and harmful cells from the blood into the nervous tissue (Wilhelm et al., 2013). Before being able to move through the BBB, cancer cells spend several days stalled in the capillary lumen and induce several changes in the endothelium, possibly making it more permissive (Haskó et al., 2019). During this period, tumour cells may communicate with cerebral endothelial cells through diverse pathways, including extracellular vesicles (EVs).

Tumour-derived EVs were previously shown to be taken up by, and also to cross, brain endothelial cells via endocytosis and transcytosis, respectively (Morad et al., 2019). One of the main cargos of EVs is microRNAs (miRNAs), among which breast cancer-derived miR-105 and miR-181c have been shown to compromise the integrity of the BBB by disrupting interendothelial tight junctions (TJs), thereby promoting migration of tumour cells through the walls of cerebral capillaries and the development of brain metastases (Tominaga et al., 2015; Zhou et al., 2014). However, the role of miRNAs released by breast cancer cells in the formation of brain metastases is far from being comprehensively characterized (Sereno et al., 2020).

Moreover, EVs may carry numerous other active molecules, both within their vesicular lumen and on their membranes. Transforming growth factor-β1 (TGF-β1) is one example – a cytokine released in large amounts by tumour cells, mainly via EVs (Hosseini et al., 2023). TGF-β1 regulates various processes in the tumour microenvironment, including the endothelial– mesenchymal transition of brain endothelial cells, resulting in compromised TJs and enhanced transendothelial migration of tumour cells (Krizbai et al., 2015).

Therefore, our main goal was to identify relevant miRNAs and other physiologically active substances carried by the EVs of brain metastatic TNBCs that can open the BBB and consequently enhance migration of the tumour cells into the brain. In addition, we aimed to describe the molecular mechanisms underlying these phenomena and to identify possible therapeutic or secondary prevention targets in brain metastatic disease.

## Material and methods

### Collection and characterization of 4T1-tdTomato-derived EVs

The 4T1-tdTomato cell line was generated by transfection of 4T1 murine TNBC cells with the pcDNA3.1(+)/Luc2-tdTomato plasmid to express the tdTomato red fluorescent protein (RFP) (Haskó et al., 2019). Cells were maintained in RPMI-1640 medium supplemented with 5% foetal bovine serum (FBS) and geneticin (G418) for selection (all from Thermo Fisher Scientific, Waltham, MA USA). The cells were authenticated by analysis of highly variable short tandem repeat (STR) loci (Microsynth, Balgach, Switzerland) and regularly verified for the absence of mycoplasma contamination using the MycoAlert kit (Lonza, Basel, Switzerland).

EV collection and handling were performed in accordance with current guidelines (Welsh et al., 2024). For EV isolation, cells were plated into 10 cm diameter culture dishes (TPP Techno Plastic Products, Trasadingen, Switzerland). At 70–80% confluence, the cells were washed twice with phosphate-buffered saline (PBS) and supplied with fresh medium containing 2% exosome-depleted FBS (Thermo Fisher Scientific). After 24 hours, the medium was collected, and EVs were isolated using the ultracentrifugation method. Briefly, following three consecutive centrifugation steps of the supernatants (700 × g for 5 minutes, 1000 × g for 8 minutes, and 10,000 × g for 30 minutes using a Sorvall RC6+ centrifuge (Thermo Fisher Scientific), the supernatant was filtered through a 0.22 μm pore-size membrane (Merck/MilliporeSigma, Burlington, MA USA) followed by centrifugation at 150,000 × g for 90 minutes using a Sorvall wX+ultraseries centrifuge and a T1270 fixed angle rotor, all performed at 4°C. EVs were collected in PBS and stored at –20°C for maximum 3 months.

For PKH26 staining (Merck/MilliporeSigma), the volumes of EVs or vehicle PBS controls were adjusted to 0.5 ml using Diluent C. PKH26 dye was prepared by diluting 4 µl of dye in 1 ml of Diluent C and 0.5 ml of this solution was added to each sample. Samples were incubated for 5 minutes at room temperature. The labelling reaction was quenched by transferring samples to new ultracentrifuge tubes and adding 1 ml of 1% bovine serum albumin (BSA) in PBS (prefiltered through a 0.1 µm pore-size membrane), followed by gentle mixing. The volume was then adjusted to 8 ml with PBS, and samples were ultracentrifuged at 150,000 × g for 90 minutes at 4°C.

The total protein concentration of EVs was determined with MicroBCA (Thermo Fisher Scientific), while their size and morphology were characterized using dynamic light scattering and atomic force microscopy, as described in **Suppl. methods**. The EVs had an average size of 100–200 nm and exhibited the expected spherical morphology **(Suppl. fig. 1 A, B)**. They expressed CD81, Hsp70, and Alix, but not Grp94, as assessed by western blot **(Suppl. fig. 1 C)**.

### Isolation, culture, and treatments of mouse brain endothelial cells (MBECs)

We used freshly isolated MBECs in our experiments because of their superior barrier properties compared to cell lines. The cells were isolated from 6–8-week-old FVB/Ant:TgCAG-YFP mice (kindly provided by the Institute of Experimental Medicine, Budapest, Hungary), which express Venus-YFP in endothelial cells, using an established protocol. Briefly, brains were collected under sterile conditions, the meninges were removed, and the tissue was enzymatically dissociated in two steps using collagenase 2 and collagenase/dispase (Merck/MilliporeSigma). Myelin was removed by centrifugation through 20% BSA. Microvascular fragments were enriched by density separation on a 33% Percoll (Merck/MilliporeSigma) gradient. The resulting endothelial cell fraction was seeded onto fibronectin- and collagen IV-coated culture surfaces and maintained in DMEM/F12 medium containing 10% plasma-derived serum (PDS; First Link, Wolverhampton, UK), along with heparin and growth factors, as well as puromycin for initial selection. MBECs were used at passage number P1, except for RNA transfections which were performed at P0.

EVs were added to confluent MBECs cultured in 3.5 cm diameter dishes in a dose of 35 μg EV-protein/ml/dish in a modified culture media containing 5% EV-depleted PDS, in the absence of heparin. Depletion was performed by seeding of the EVs using the ultracentrifugation method.

Transfection of P0 MBECs with the synthetic miR-146a-MIMIC (Qiagen, Hilden, Germany) or *Paqr5* stealth siRNA (Thermo Fisher Scientific) was performed in 3.5 cm diameter culture dishes in serum-free OptiMEM using 2.5 μl Lipofectamine 3000 (Thermo Fisher Scientific) and 5 nM RNA. RNA sequences are indicated in **Suppl. table 1**. Transfections were carried out at 50–60% cell confluence on two successive days for 6–8 hours each, after which the cultures were allowed to recover for 24 hours before passing the cells for subsequent experiments.

All treatments were performed at 100% confluence. TGF-β1 (PeproTech, Cranbury, NJ USA) was applied on the cells in a concentration of 10 ng/ml, while SB-431542 (Merck/MilliporeSigma) was used in a final dose of 10 μM for 24 hours in the modified culture media containing 5% EV-depleted PDS.

### RNA extraction, NanoString nCounter analysis, cDNA synthesis, and qPCR

Total RNA was extracted using the miRNeasy kit (Qiagen) according to the manufacturer’s instructions. RNA concentration was measured with a NanoDrop (Thermo Fisher Scientific) or a DS-11 FX+ spectrofluorimeter.

For screening of miRNAs, the nCounter miRNA Expression Assay and the mouse v1.5 miRNA codeset (NanoString Technologies, Seattle, WA USA) were used, covering 577 mouse miRNAs. For the profiling, 100 ng total RNA/sample was used from the following samples: 4T1-tdTomato cells, 4T1-tdTomato EVs, control MBECs, and MBECs receiving 35 μg EV-protein/ml/dish EVs for 24 hours. Assessment of RNA quality using the Agilent Bioanalyzer system, ligation of miRtags, hybridisation to probes, washing, immobilisation to the solid substrate, scanning, and data collection were performed by RT-Europe Research Center (Mosonmagyaróvár, Hungary). Raw count data obtained from the nCounter analysis were normalized to positive controls and processed to obtain a list of potential changed miRNA. As a background threshold, the mean of 8 negative controls + 3× SD was applied, and only the counts above the threshold were taken into consideration. The geometric mean of three housekeeping genes (*Actb, B2m*, and *Rpl19*) was used for data normalization. Two independent experiments were performed.

For qPCR analysis of miRNAs, 200 ng RNA was transcribed into cDNA, using the miRCURY LNA RT kit (Qiagen). As a quality control for cDNA synthesis, the synthetic RNA Spike-in Uni-SP6 (Qiagen) was added to the mixture. For the qPCR reaction, the miRCURY LNA SYBR Green PCR kit (Qiagen) was used according to manufacturer’s instructions, using a cDNA dilution of 1:4. The following conditions were used: 50 cycles of 95°C for 15 seconds, 56°C for 30 seconds, and 72°C for 30 seconds. Predesigned LNA primer pairs were purchased from Qiagen for the selected miRNAs, as shown in **Suppl. table 2**.

For reverse transcription of mRNAs, 1 μg RNA and the Maxima first strand cDNA synthesis kit (Thermo Fisher Scientific) was used. The qPCR reaction was performed using Luminaris color HiGreen qPCR master mix (Thermo Fisher Scientific). The protocol consisted of 40 cycles of 95°C for 15 seconds, 60°C or 64°C for 30 seconds and 72°C for 30 seconds. Primers were designed with the NCBI Primer Blast tool and the sequences are listed in **Suppl. table 3**.

All PCR reactions were performed on an iQ5 or a CFX96 thermocycler (Bio-Rad, Hercules, CA USA) in 96-well plates, in triplicates and a no-template control was included for each amplification. Threshold cycles were determined using the software of the thermocycler, while the quantitation was determined with the ΔΔCt method.

### Prediction of miR-146a-5p mRNA targets

To identify potential targets of mmu-miR-146a-5p, a comparative bioinformatics analysis was performed using four online prediction tools. First, miRDB (Chen and Wang, 2020) was used to obtain support vector machine-based predictions and targets with scores between 80 and 100 were considered reliable. DIANA microT-CDS predictions (Reczko et al., 2012) were evaluated using the recommended threshold of 0.7. Results from TargetScan (McGeary et al., 2019) were ranked according to the cumulative weighted context++ score, total context score, and the aggregate probability of conserved targeting. Finally, miRmap, an open-source framework integrating thermodynamic, evolutionary, and probabilistic features, was additionally used to rank targets based on predicted repression strength expressed as the miRmap score (Vejnar et al., 2013). Filters applied in this analysis included ΔG and miRmap score thresholds of 70.

To refine the candidate list, only genes predicted by at least three of the four tools were retained. Given the large number of potential targets, the results were further cross-referenced with the Betsholtz database (https://betsholtzlab.org/VascularSingleCells/database.html) to identify transcripts expressed in brain endothelial cells and to prioritize those most relevant for subsequent validation.

### *In vivo* brain metastasis model

Young adult (8-week-old, 20-22 g) female FVB/Ant:TgCAG-YFP mice were housed under standard conditions (12-hour light/dark cycle, 23±2°C) with *ad libitum* access to regular chow and water.

Brain metastases were generated as reported previously (Haskó et al., 2019). Briefly, under isoflurane anaesthesia, mice were inoculated with 10^6^ 4T1-tdTomato cells suspended in 200 µl of sterile Krebs–Ringer solution into the right common carotid artery using a 30G cannula, while the right external carotid artery was transiently ligated. The wound was closed and the animals were let to recover. 48 hours later, mice were anaesthetised with Avertin, perfused with saline and 4% paraformaldehyde (PFA) and processed further for either *in situ* hybridization or immunofluorescence studies.

### RNAscope *in situ* hybridization

Brains were collected, post-fixed in 4% PFA overnight, and subsequently cryoprotected in 30% sucrose in PBS for 72 hours. The tissues were then embedded in Tissue-Tek OCT compound (Sakura Finetek, Tokyo, Japan) and frozen using an isopentane/dry-ice cooling method. Coronal brain sections (15 µm) were prepared using a cryostat (Leica, Wetzlar, Germany). Six different cortical and mid-brain regions were selected for downstream analyses. *In situ* hybridization and immunofluorescence were performed in a sequential workflow. *In situ* hybridization was performed with the RNAscope Multiplex Fluorescent Reagent Kit v.2 combined with the RNA-Protein Co-detection Ancillary Kit (Bio-Techne, Minneapolis, MI USA) using the Mm-Paqr5-C1 probe (cat no. 1066291-C1) and appropriate negative and positive controls. Labelling was performed with the TSA Vivid Fluorophore Kit 650. Because the fluorescence intensity of fluorescent proteins may decrease during the *in situ* hybridization protocol, immunofluorescence labelling was performed to visualize endothelial and tumour cells. Samples were treated with the co-detection blocker and incubated overnight at 4°C with primary antibodies against CD31 and RFP (**Suppl. table 4**). Following three washing steps with PBS, Alexa Fluor 488- and Alexa Fluor 594-conjugated secondary antibodies (**Suppl. table 5**) were added to the samples. After washing, DAPI (Bio-Techne, Minneapolis, MI USA) was used to counterstain cell nuclei and Fluoromount-G (Thermo Fisher Scientific) was applied to mount the samples.

Imaging was conducted with a 60× oil-immersion objective on a VisiScope spinning-disk confocal microscope (Visitron Systems, Puchheim, Germany). Images were exported in Tagged Image File (TIF) format, after which a custom made MATLAB (MathWorks, Natick, MA USA) routine was used to quantify the specific staining spots (dots). Stacked images were treated as three-dimensional structures during analysis. Image volumes were segmented using two thresholding criteria. First, a size-based threshold was applied to exclude undersized detections (<50 voxels, corresponding to approximately 1 μm^3^) as well as large clusters exceeding 2.5 times the size of a typical detected dot, which were considered indistinguishable from potential imaging artefacts. Second, an intensity-based threshold was applied, retaining only dots with mean intensities within the upper quartile of the image brightness histogram. Notably, the intensity-based exclusion was substantially more stringent than the size-based exclusion. For vessel and tumour volume quantification, all voxels with intensities above one-third of the intensity histogram of the respective detection channel were included. Dots across channels were considered co-localized (positive hits) if they overlapped or were located within a three-voxel vicinity of the vessel or tumour volumes. Specificity and scoring accuracy were validated by comparing machine-generated counts with manual counts across five images, with discrepancies remaining below 5%. Data from three animals were included in the statistical analysis.

### Immunofluorescence staining of cell cultures and brain sections

Cell cultures were fixed with 4% PFA for 10 minutes at room temperature, washed afterwards with PBS, and permeabilized with 0.2% Triton X-100 in PBS for 15 minutes. After blocking with 3% BSA (Merck/MilliporeSigma) for 20 minutes, samples were incubated with primary antibodies against ZO-1 or claudin-5 (**Suppl. table 4**). The staining was visualized using an Alexa Fluor 647-conjugated secondary antibody (**Suppl. table 5**). Nuclei were stained with Hoechst 33342 (Merck/MilliporeSigma). Samples were mounted in Fluoromount-G and studied with an Eclipse TE2000 U inverted microscope (Nikon, Tokyo, Japan).

For the detection of Paqr5/mPRγ in brain sections, 20 µm-thick floating sections were prepared using a freezing microtome (Reichert-Jung, Leica). All sections were rinsed in 0.5% Tween-20 in PBS, then blocked for 30 minutes in a blocking buffer containing 2% donkey serum (Merck/MilliporeSigma), 50 mM glycine, 0.1% Triton X-100, 0.1% BSA, and 0.05% Tween-20 in PBS (Flores-Maldonado et al., 2020). The primary antibody (**Suppl. table 4**) was diluted in a solution containing 10 mM glycine, 0.1% Triton X-100, and 0.05 % Tween-20 in PBS and sections were incubated overnight at 4°C on an orbital shaker. Slides were extensively washed in 0.5% Tween-20 in PBS, and an Alexa Fluor 647-labelled secondary antibody was applied in 0.1% Tween-20 in PBS for two hours at room temperature in the dark. Sections were washed, counterstained with Hoechst 33342 for 10 minutes, washed again with PBS, rinsed in water, and mounted in FluoroMount-G. Samples were analysed using a VisiScope spinning-disk confocal microscope.

### Assessment of BBB functions *in vitro*

For impedance/cell index measurements, cells were seeded at a density of 75,000 cells/cm^2^ on E-plates having golden electrodes in the bottom of the wells. Impedance of the cell layers was measured with the xCELLigence RTCA instrument (ACEA Biosciences, San Diego, CA USA).

For transendothelial electric resistance (TEER) measurements, MBECs were seeded at a density of 75,000 cells/cm^2^ on semipermeable filter inserts with 0.4 μm pore size (Corning-Costar, Corning, NY USA). After reaching confluence, the monolayers were supplied with 550 nM hydrocortisone, 250 μM CPT-cAMP, and 17.5 μM Ro-20-1724 (all from Merck/MilliporeSigma) to induce maturation of TJs. TEER was measured with the cellZscope instrument (nanoAnalytics, Münster, Germany). After completing the TEER measurements, sodium fluorescein permeability was assessed in the same cells.

After removal from the cellZscope instrument, filters were washed with serum-free and phenol red-free DMEM. Sodium fluorescein was applied in the top compartment in a final concentration of 10 μg/ml. The bottom compartment was loaded with phenol red-free DMEM. Filters were placed to 37°C and gentle shaking (110 rpm) was applied. Samples were taken from the basolateral side after 30 minutes and the analysis was performed with a FLUOstar Optima fluorescent microplate reader (BMG Labtechnologies, Offenburg, Germany). The apparent permeability was calculated using the formula: Papp = dQ/(dT × A × C_0_), where dQ is the transported amount, dT is the incubation time, A is the surface of filter (0.33 cm^2^), and C_0_ is the initial concentration.

### Western blot

Cells were lysed in ice-cold buffer (20⍰mM Tris, 150⍰mM NaCl, 0.5% Triton X-100, 1% sodium deoxycholate, 0.1% SDS, 1⍰mM sodium orthovanadate, 10⍰mM sodium fluoride, 1⍰mM EDTA, and 1⍰mM Pefabloc) and incubated on ice for 30⍰minutes, followed by centrifugation at 10,000 × g for 10⍰minutes at 4°C. Protein concentration was measured with BCA (Thermo Fisher Scientific) using the DS-11 FX+ spectrofluorimeter (DeNovix, Wilmington, DE USA). Laemmli buffer was added and samples were heated at 60°C for 3 minutes. Proteins were separated by SDS-PAGE and transferred onto 0.2⍰μm pore-size PVDF or nitrocellulose membranes (Bio-Rad). Membranes were blocked with 3% BSA in Tris-buffered saline with 0.1% Tween-20 (TBS-T) for 60⍰minutes at room temperature and then incubated overnight at 4°C with primary antibodies (**Suppl. table 4**) in TBS-T. After washing, membranes were incubated with the appropriate secondary antibodies (**Suppl. table 5**) for 1 hour at room temperature, followed by additional washes. Immunoreactive bands were visualized using the Clarity Enhanced Chemiluminescence Substrate (Bio-Rad) on a ChemiDoc MP System (Bio-Rad) and band intensities were quantified with Image Lab software v5.2 (Bio-Rad) and normalized to β-actin.

### Transmigration assay

Transmigration experiments were performed as described previously with slight modifications (Wilhelm et al., 2014). Briefly, MBECs were passed onto 8 μm pore-size filter inserts (Corning) having fibronectin and collagen IV coating on the top side and Growth Factor-Reduced Matrigel (Corning) on the bottom side, preventing endothelial cells from dropping into the pores. After reaching confluence, MBECs were supplemented with hydrocortisone, CPT-cAMP, and Ro 20-1724 from the top side and primary mouse astrocyte-conditioned medium from the basolateral side. After 8 hours, 10^5^ 4T1-tdTomato cells/cm^2^ were plated onto the MBEC cell layer. Because breast cancer cells are less effective at opening the BBB than melanoma cells, the transmigration assay was performed for a longer duration than the 5 hours used for the latter, while remaining short enough to preserve the integrity of the endothelial monolayer in the absence of tumour cells. Accordingly, 4T1-tdTomato cells were incubated on MBECs for 48 hours, followed by fixation with 4% PFA. Cells from the top side of the filters were removed with a cotton swab and breast cancer cells migrated through the endothelial monolayer and the pores of the filter were imaged with an Axio Observer Z1 inverted epifluorescence microscope (Zeiss, Oberkochen, Germany) equipped with a STEDYCON laser scanning confocal microscope (Abberior Instruments, Göttingen, Germany). A custom-made MATLAB routine was used to threshold segmented images and quantify transmigrated cells, defined as structures with the expected morphology and size.

## Results

### EVs released by TNBC cells increase miR-146a-5p levels in brain endothelial cells

To investigate how tumour cell-derived EVs affect BBB-forming endothelial cells during pre-metastatic niche formation, we exposed primary MBECs to EVs isolated from 4T1-tdTomato culture supernatants. When PKH26-labelled EVs were added to MBEC cultures, we observed that endothelial cells internalized the vesicles, which accumulated in the perinuclear region. In parallel, TJs exhibited minor alterations, reflected by a disorganized localization of ZO-1, a protein of the TJ cytoplasmic plaque (**Fig. 1 A**).

**Figure 1.**
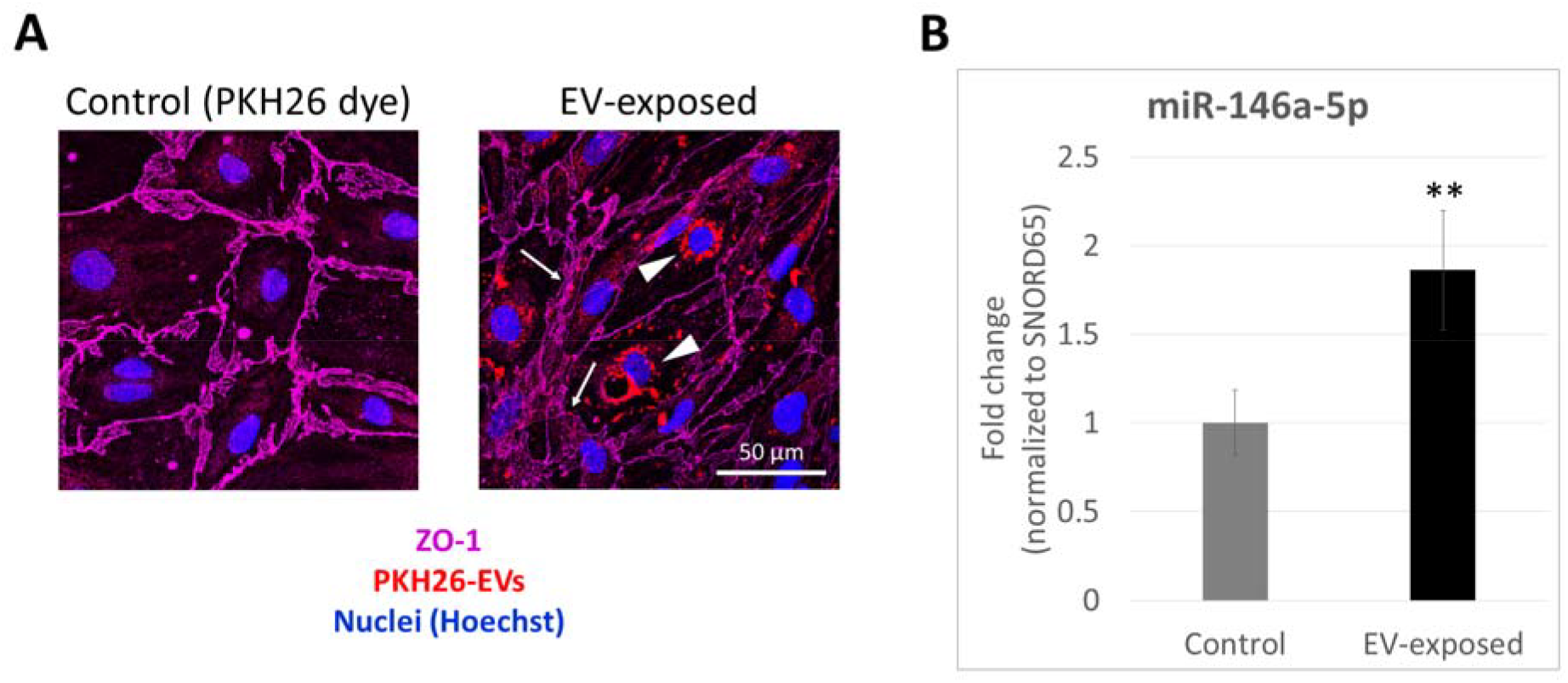
Uptake of TNBC cell-derived EVs and the subsequent elevation of miR-146a-5p levels in cerebral endothelial cells. **A:** Confluent MBEC monolayers were exposed to PKH26-labelled 4T1-tdTomato-derived EVs in a dose of 35 μg EV-protein/ml, while control cells received the respective amount of the PKH26 dye. After 24 hours, cells were fixed and stained for ZO-1. Representative confocal micrographs (maximum intensity projection of *z*-stacks) are shown from N = 3 independent experiments. Arrows indicate disorganized junctions, while arrowheads show perinuclear EVs. **B:** MBECs were exposed to 4T1-tdTomato-derived EVs in a dose of 35 μg EV-protein/ml. After 24 hours, total RNA was isolated and miR-146a-5p levels were determined by qPCR. Graph represents average ± SD from N = 4 independent experiments, each performed in triplicates. **P ≤ 0.01 compared to control (Student’s *t*-test).

MiRNAs represent a major cargo of EVs. To identify those changed in cerebral endothelial cells in response to tumour-derived EVs, we performed a NanoString nCounter screening assay. By comparing control and EV-exposed MBECs and applying stringent selection criteria – including high expression of the respective miRNA in both tumour cells and EVs, high read counts in the samples, and a change of more than 1.5-fold in two independent experiments – the amount of miR-146a-5p was found to be increased in MBECs receiving the EVs. Using qPCR, we validated the higher miR-146a-5p count in MBECs encountering tumour-released EVs (**Fig. 1 B**).

### *Paqr5* is downregulated by miR-146a-5p in brain endothelial cells

Knowing that miRNAs usually induce degradation or translational repression of certain specific mRNAs (Naeli et al., 2023), we next aimed to identify the target genes of miR-146a-5p. For this, we used four different bioinformatics tools (miRDB, DIANA microT-CDS, TargetScan, and miRmap) and selected those mRNAs, which were predicted by at least three algorithms to be possibly regulated by miR-146a-5p. Using this method, five genes were identified: *Med1* (encoding MED1, mediator complex subunit 1), *Numb* (translating to NUMB), *Traf6* (the gene of TRAF6, tumour necrosis factor receptor-associated factor 6), *Irak1* (encoding IRAK1, interleukin-1 receptor-associated kinase 1), and *Paqr5* (translating to PAQR5, the progestin and adipoQ receptor family member 5 or membrane progesterone receptor γ (mPRγ) protein). We confirmed the downregulation of all five genes in MBECs exposed to EVs using qPCR (**Fig. 2 A**).

**Figure 2.**
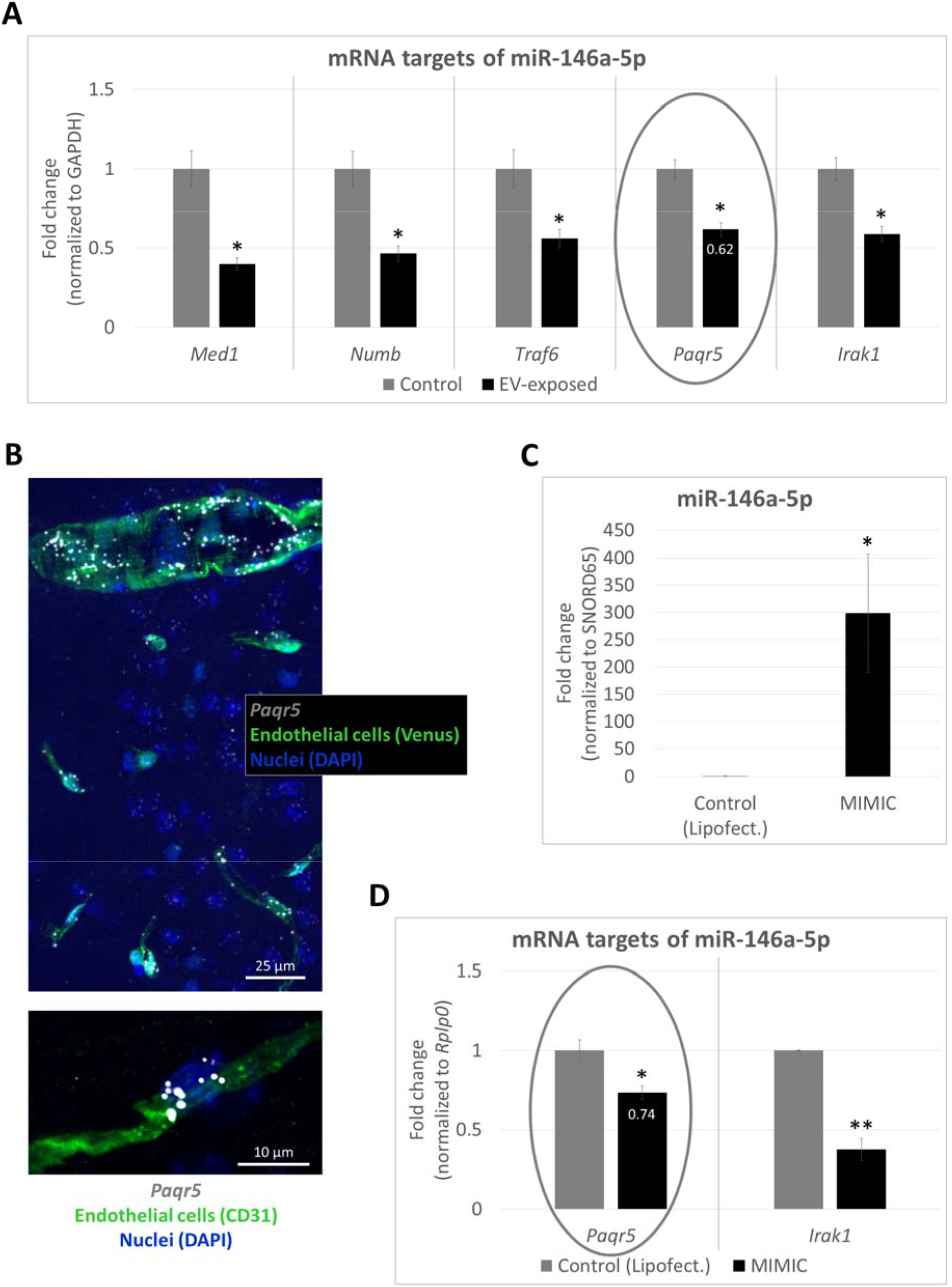
Targets of miR-146a-5p. **A:** MBECs were exposed to 4T1-tdTomato-derived EVs in a dose of 35 μg EV-protein/ml. After 24 hours, total RNA was isolated and levels of the mRNA targets were determined by qPCR. Graph represents average ± SD from N = 2 independent experiments, each performed in triplicates. *P ≤ 0.05 compared to control (Student’s *t*-test for each gene individually). **B:** Expression of *Paqr5* was determined by RNAscope *in situ* hybridization in mouse brain sections. Representative confocal micrographs (maximum intensity projection of *z*-stacks) are shown from N = 3 animals. **C, D:** MBECs were transfected twice with 5 nM miR-146-MIMIC using Lipofectamine, while control cells received the transfection agent only. 24 hours after removal of the transfection reagents, total RNA was isolated and miR-146a-5p, *Paqr5*, and *Irak1* levels were determined by qPCR. Graphs represent average ± SD from N = 2 independent experiments, each performed in triplicates. *P ≤ 0.05, **P ≤ 0.01 compared to Lipofectamine control (Student’s *t*-test for each RNA individually).

Among the five selected genes, four – *Med1, Numb, Traf6*, and *Irak1* – are well-known elements of certain pathways regulating cell division, inflammation, and tumourigenesis and are widely expressed in diverse cell types. On the other hand, the functions of *Paqr5* are largely uncharacterized, while its expression is mostly restricted to endothelial cells in the brain. This was indicated by single cell RNA-sequencing results of brain vascular cells (https://betsholtzlab.org/VascularSingleCells/database.html (Vanlandewijck et al., 2018)) (**Suppl. fig. 2 A**) and data from The Allen Brain Cell Atlas as well (https://portal.brain-map.org/atlases-and-data/bkp/abc-atlas (Yao et al., 2023)) (not shown). Our *in situ* hybridization data confirmed endothelial-specific localization of *Paqr5* both in large and small vessels (**Fig. 2 B**) without differences in adult females and males (not shown). Selective endothelial expression is a feature of only *Paqr5* among members of the mPR family, as *Paqr6, Paqr7*, and *Paqr8* are much less abundant in endothelial cells of the brain than *Paqr5*. In addition, *Paqr6, Paqr7*, and *Paqr8* all show higher expression in astrocytes than in cerebral endothelial cells, as indicated by RNA sequencing data of The Allen Brain Cell Atlas (not shown) and our qPCR results as well (**Suppl. fig. 2 B**).

To confirm experimentally that miR-146a is responsible for *Paqr5* downregulation in brain endothelial cells exposed to EVs, we transfected the cells with a synthetic miR-146a-MIMIC resulting in a ~300-fold increase in the miR-146a-5p content of the cells (**Fig. 2 C**). Decreased expression of *Paqr5* was indeed detected in these cells; however, this was lower than that induced by the EVs (0.74 *vs*. 0.62 fold change) (**Fig. 2 D**) despite the much higher increase in the miR-146a-5p levels. On the other hand, reduction in the *Irak1* mRNA levels – one of the best-known targets of miR-146a – was more pronounced in the MIMIC-transfected cells than in EV-treated cells (**Fig. 2 A** and **D**). This suggested that another mechanism might be also involved in EV-related *Paqr5*, but not *Irak1* downregulation in brain endothelial cells, which is independent of miR-146-5p.

### TGF-β1 and miR-146a-5p collectively regulate *Paqr5* in brain endothelial cells exposed to tumour-derived EVs

The main negative regulator of PAQR5 is TGF-β1, which has been previously shown to suppress *Paqr5* expression in renal carcinoma cells (Tao et al., 2022). Moreover, TGF-β1 is a major protein cargo of tumour-released EVs (Hosseini et al., 2023). Therefore, we tested the effect of TGF-β1 on the expression of the five miR-146a-5p-repressed genes. For inhibition of type I TGF-β receptors SB-431542 was used. No change in the expression of *Numb, Traf6*, and *Irak1* was detected in response to TGF-β1 or TGF-β1 + SB-431542 (**Fig. 3 A**). On the other hand, TGF-β1 induced an increase in *Med1* expression, which was partly abolished by the receptor inhibitor. This suggests a delicate interplay between TGF-β1 and MED1 signalling in endothelial cells, as previously MED1 was shown to control the TGF-β axis in the pulmonary arterial endothelium (Wang et al., 2022).

**Figure 3.**
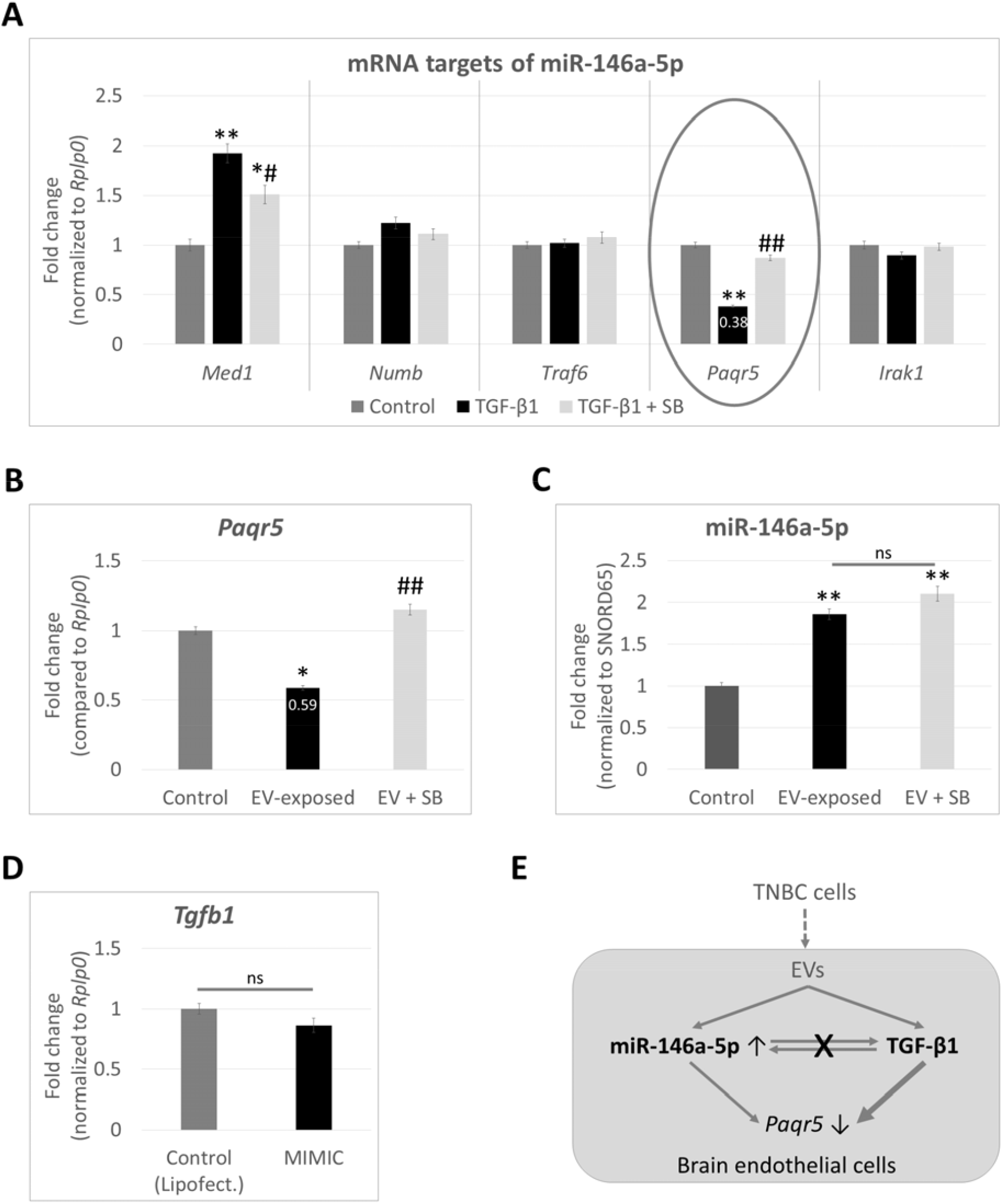
Regulation of *Paqr5* by miR-146a-5p and TGF-β1. **A**: MBECs were exposed to 10 ng/ml TGF-β1 in the absence or presence of SB-431542, a type I TGF-β receptor inhibitor. After 24 hours, total RNA was isolated and levels of the mRNA targets of miR-146a-5p were determined by qPCR. Graph represents average ± SD from N = 2 independent experiments, each performed in triplicates. * P ≤ 0.05, * * P ≤ 0.01 compared to control, ^#^P ≤ 0.05, ^##^P ≤ 0.01 compared to TGF-β1-treated cells (ANOVA and Tukey *post-hoc* test for each gene individually). **B, C:** MBECs were exposed to 4T1-tdTomato-derived EVs in a dose of 35 μg EV-protein/ml, in the absence or presence of SB-431542. After 24 hours, total RNA was isolated and *Paqr5* or miR-146a-5p levels were determined by qPCR. Graphs represent average ± SD from N = 2 independent experiments, performed in triplicates. * P ≤ 0.05, * * P ≤ 0.01 compared to control, ^##^P ≤ 0.01 compared to EV-treated cells, ns = not significant (ANOVA and Tukey *post-hoc* test). **D:** MBECs were transfected twice with 5 nM miR-146-MIMIC using Lipofectamine, while control cells received the transfection agent only. 24 hours after removal of the transfection reagents, total RNA was isolated and *Tgfb1* mRNA levels were determined by qPCR. Graph represents average ± SD from N = 2 independent experiments, each performed in triplicates. ns = not significant (Student’s *t*-test). E: The cartoon is a schematic representation of the two parallel mechanisms through which TNBC-derived EVs downregulate *Paqr5* in brain endothelial cells during early phases of metastasis formation.

More importantly, TGF-β1 downregulated *Paqr5* expression in brain endothelial cells. This effect was more potent than that of miR-146a-5p (0.38 *vs*. 0.74 fold change) (**Fig 2 D** and **Fig. 3 A**) and mediated through type I TGF-β receptors. In addition, SB-431542 reversed the EV-induced reduction in *Paqr5* expression (**Fig. 3 B**), confirming that TGF-β1 is a more potent suppressor of *Paqr5* than miR-146a-5p. Importantly, SB-431542 did not alter the amount miR-146a-5p in EV-exposed brain endothelial cells (**Fig. 3 C**), while transfection of the synthetic miR-146a-MIMIC into these cells did not change their *Tgfb1* levels (**Fig. 3 D**). These results indicate that TGF-β1 is neither upstream nor downstream of miR-146a-5p in this pathway.

Altogether, our data suggest that TNBC cell-released EVs induce repression of *Paqr5* expression in brain endothelial cells through two mechanisms: miR-146a-5p and, more potently, TGF-β1 (**Fig. 3 E**).

Although the differences were less pronounced than in mouse cells, similar changes were detected in D3 human brain endothelial cells exposed to EVs collected from the supernatant of the brain metastatic variant of MDA-MB-231 TNBC cells, called BrM2. In response to BrM2-derived EVs, we detected increase in the miR-146a-5p and decrease in the PAQR5 mRNA content of D3 cells (**Suppl. fig. 3 A, B**). In addition, PAQR5 mRNA expression decreased upon transfection with the miR-146a-MIMIC (**Suppl. fig. 3 C, D**) and in response to TGF-β1 as well (**Suppl. fig. 3 E**). These data suggest a species-independent effect of TNBC cell-released EVs on brain endothelial cells, namely a miR-146a-5p- and TGF-β1-induced downregulation of *Paqr5*/PAQR5 in both mouse and human endothelial cells of the BBB.

### *Paqr5* is downregulated in brain vessels containing arrested tumour cells in mice

Next, we sought to determine whether metastatic tumour cells also downregulate *Paqr5* in brain vessels *in vivo*. To this end, we injected the cells into the circulation of mice having Venus-YFP-expressing endothelial cells. Based on our previous studies (Haskó et al., 2019), tumour cell transmigration occurs predominantly between days 3 and 5 after inoculation. Because our aim was to assess interactions between metastatic and cerebral endothelial cells before extravasation, we examined the brains on day 2, when the vast majority of breast cancer cells remained confined to the capillary lumen.

Using RNAscope *in situ* hybridization to detect *Paqr5* expression, we compared vessels containing tumour cells with those that were tumour-free. As shown in **Fig. 4 A**, *Paqr5* levels significantly decreased in endothelial cells coming in direct contact with arrested tumour cells.

**Figure 4.**
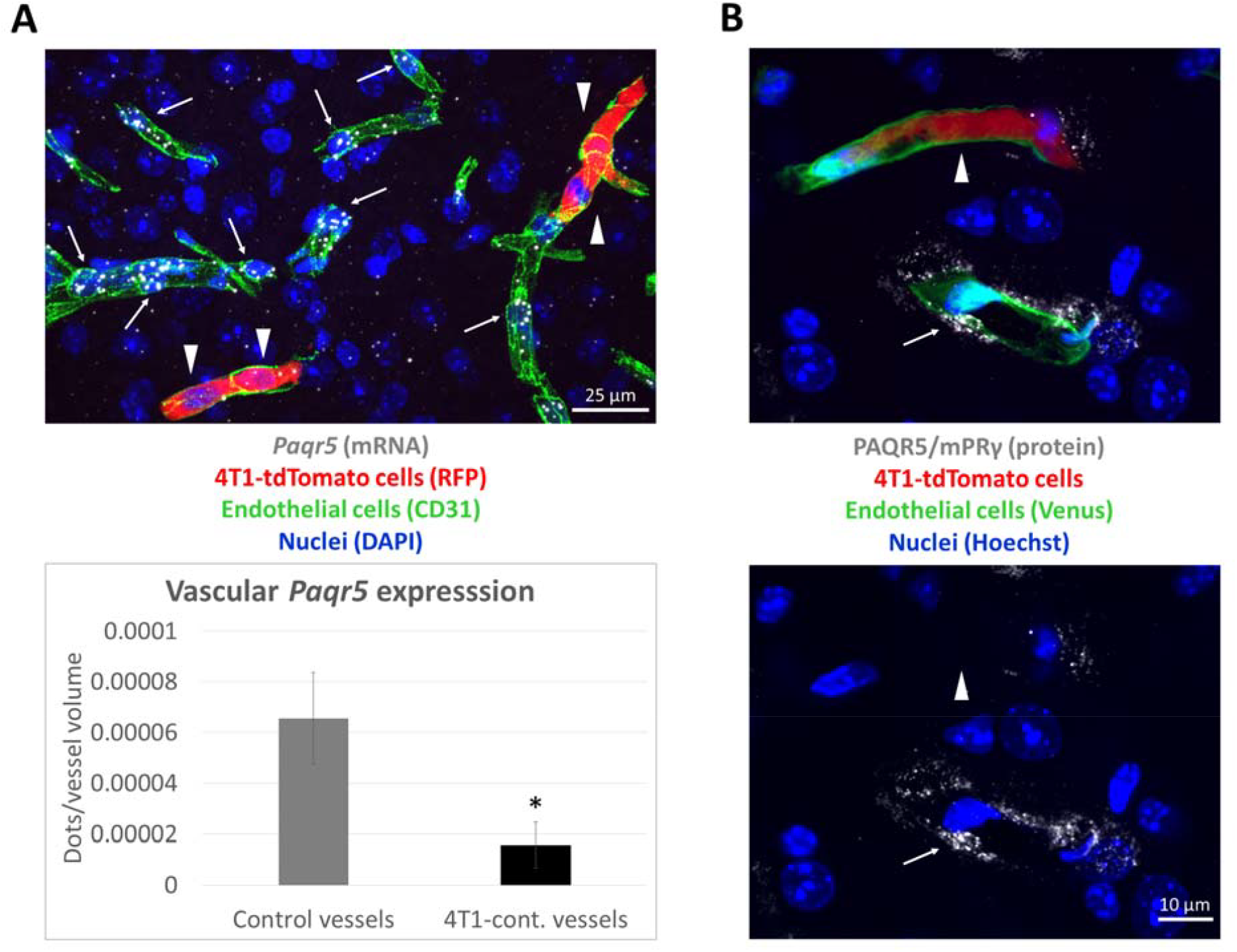
Downregulation of *Paqr5*/mPRγ expression in brain capillaries containing arrested TNBC cells. 4T1-tdTomato cells were injected into the circulation of mice through the internal carotid artery. After two days, animals were sacrificed and brain sections were prepared. **A:** *Paqr5* mRNA was detected using RNAscope *in situ* hybridization followed by immunofluorescent staining of tdTomato (RFP, red fluorescent protein) and CD31 proteins. The top panel is a representative confocal micrograph (maximum intensity projection of *z*-stacks). Graph represents quantification of *Paqr5*-positive dots in tumour-free (control) and 4T1-tdTomato cell-containing vessels. Average ± SD from N = 3 animals (n = 6 sections/mouse) is shown. *P ≤ 0.05 compared to control (Student’s *t*-test). **B:** PAQR5/mPRγ protein was detected by immunofluroescence. Representative confocal micrographs (maximum intensity projection of *z*-stacks) from a total of N = 3 animals are shown. In micrographs, arrows indicate *Paqr5*/mPRγ staining in vessels free of tumour cells, while arrowheads show the absence of *Paqr5*/mPRγ staining in brain endothelial cells coming in contact with tumour cells.

In line with the mRNA data, PAQR5/mPRγ protein levels also decreased in vessels containing arrested tumour cells (**Fig. 4 B**).

### Tumour EVs and *Paqr5* downregulation compromise barrier properties of brain endothelial cells and enhance the transendothelial migration of tumour cells

Finally, we sought to assess the functional consequences of tumour EV-induced *Paqr5* downregulation in brain endothelial cells. As key components of the BBB, these cells are characterized by continuous TJs that interconnect them into a highly impermeable vascular barrier, protecting the brain from potentially toxic substances and invading cells. Because we observed alterations in the TJs after EV uptake (**Fig. 1 A**), we conducted a more detailed analysis of the endothelial barrier integrity upon exposure to TNBC-released EVs, as well as *Paqr5* downregulation.

4T1-tdTomato EVs induced a significant decrease in the TEER of the cells, which is a direct indicator of TJ disturbance (**Fig. 5 A**). In line with this, the permeability of the EV-treated endothelial monolayers to the small molecular tracer sodium fluorescein increased significantly (**Fig. 5 B**). The backbone of endothelial TJs is formed by claudin-5, a four-transmembrane protein acting as a primary gatekeeper that restricts the passage of ions and solutes into the brain (Greene et al., 2019). Indeed, functional changes in the barrier integrity were accompanied by decreased claudin-5 levels in cells exposed to EVs (**Fig. 5 C**). On the other hand, Org OD 02-0 (10-ethenyl-19-norprogesterone, abbreviated as Org) – an mPR agonist – and progesterone as well increased claudin-5 expression in MBECs and D3 cells as well (**Suppl. fig. 4 A, B**).

**Figure 5.**
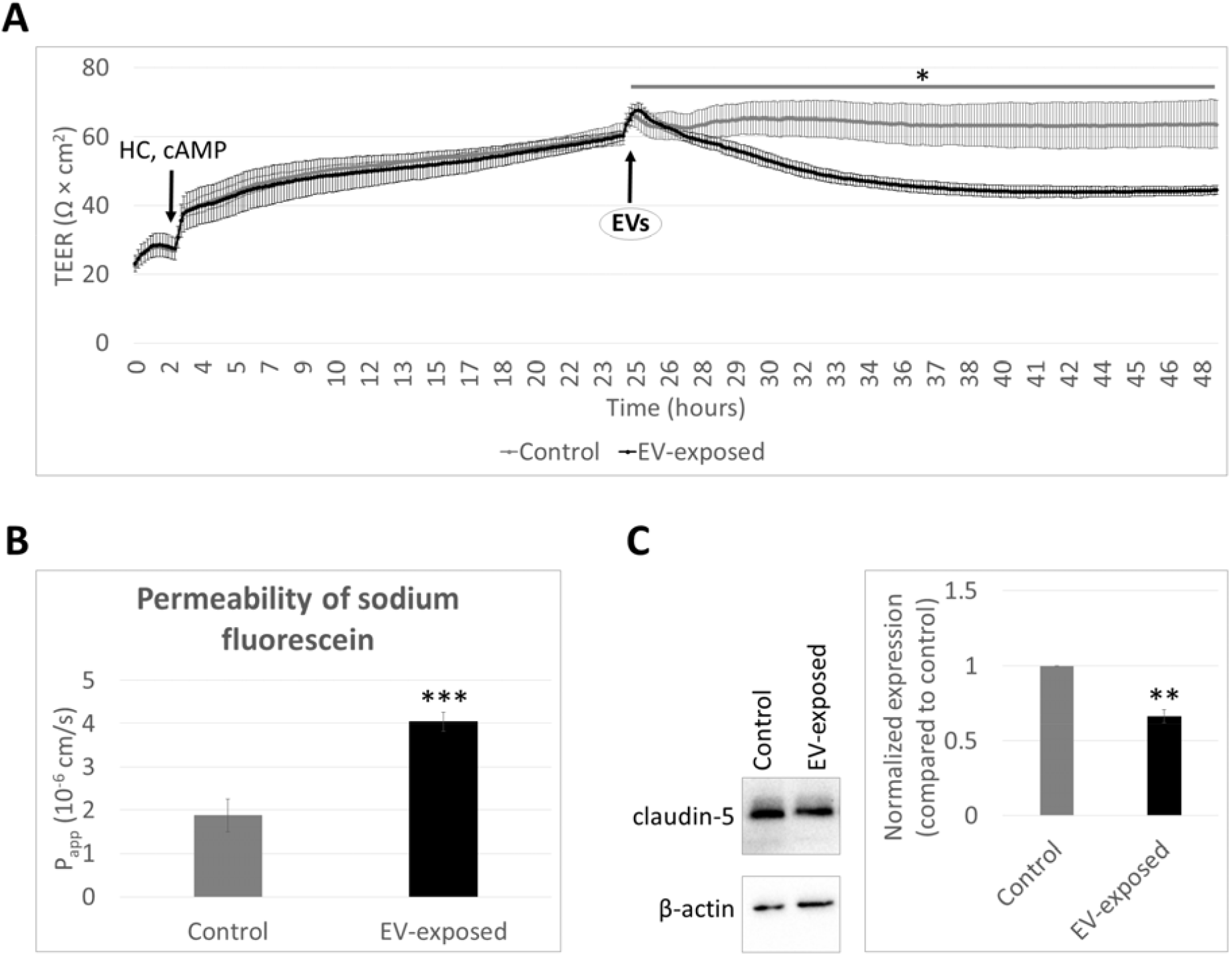
Compromised barrier properties in brain endothelial cells exposed to TNBC cell-derived EVs. **A:** MBEC monolayers cultured on semipermeable filter inserts received hydrocortisone (HC) and cAMP (CPT-cAMP + Ro 20-1724) after reaching confluence. TEER was continuously measured with the cellZscope instrument. Approximately 24 hours later, plateau was reached and the cells received 4T1-tdTomato-derived EVs in a dose of 35 μg EV-protein/ml. TEER was continuously measured for an additional 24 hours. Graph shows one representative of N = 2 independent experiments, each performed in triplicates. *P ≤ 0.05 compared to control cells (areas under curves compared with Student’s *t*-test). **B:** 24 hours after addition of the EVs, permeability of sodium fluorescein through control and EV-exposed MBEC monolayers was measured for 30 minutes. Graph shows one representative of N = 2 independent experiments, each performed in triplicates. * * *P ≤ 0.001 compared to control (Student’s *t*-test). C: Total protein was extracted for western blot from MBECs treated for 24 hours with 4T1-tdTomato-derived EVs (35 μg EV-protein/ml). Left panels show representative blots, β-actin was used as loading control. Graph represents average ± SD from N = 2 independent experiments. * *P ≤ 0.01 compared to control cells (Student’s *t*-test).

Therefore, we hypothesized, that inhibition of progesterone signalling, and especially *Paqr5* downregulation – which is induced by the tumour-derived EVs – might also compromise the barrier properties of brain endothelial cells. To clarify this question, we silenced *Paqr5* in MBECs (**Suppl. fig. 5 A**) and assessed their barrier properties. First, as a complex readout of cell layer properties – including TJ tightness, as well as cell number, viability, and adhesion – we measured the cell index of control and *Paqr5*-silenced cells cultured on golden electrodes. Using this method, we showed that *Paqr5* downregulation significantly impaired the integrity of the MBEC monolayer (**Fig. 6 A**).

**Figure 6.**
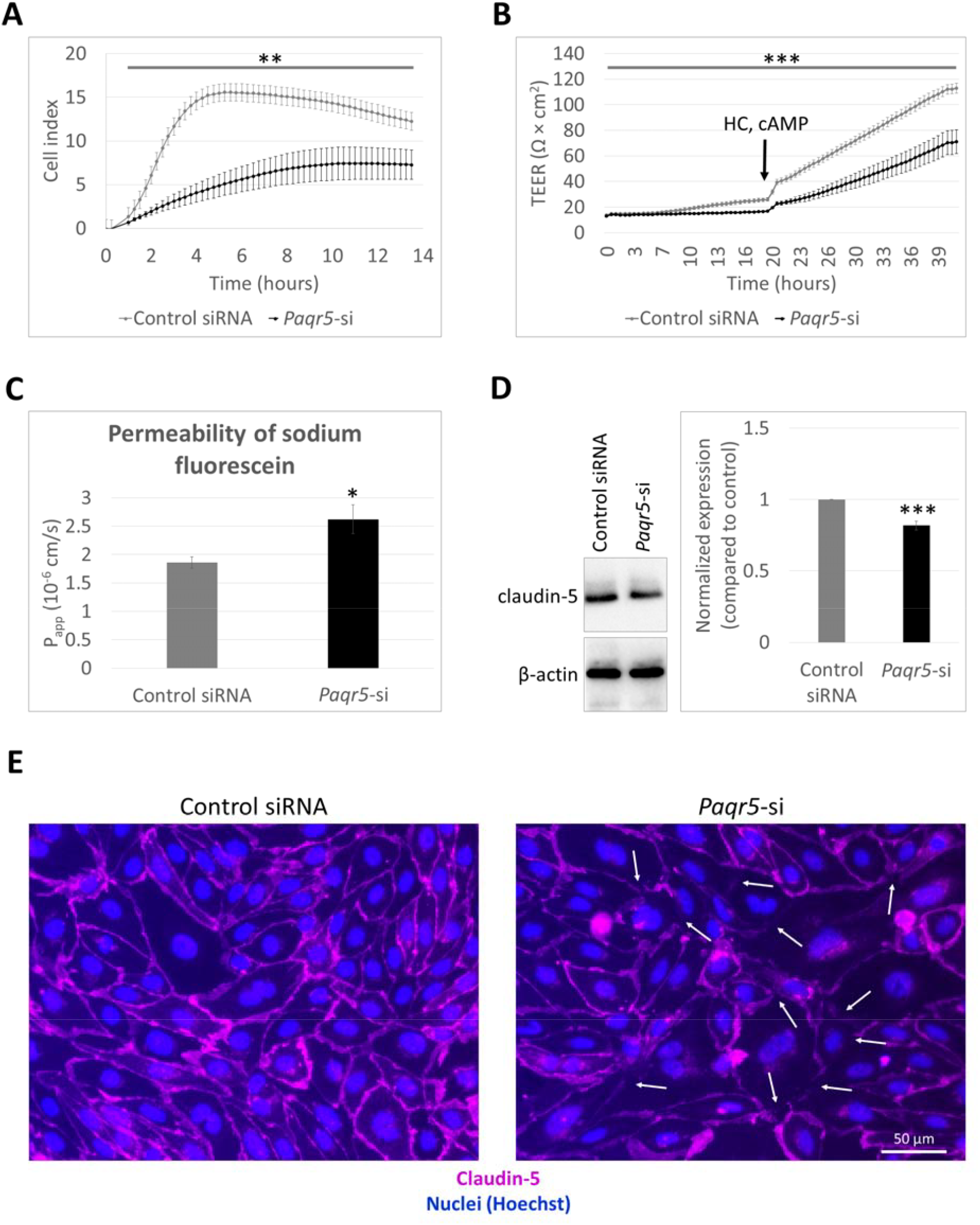
Compromised barrier properties in brain endothelial cells after silencing of *Paqr5*. Silencing of *Paqr5* in MBECs was performed with a stealth RNA construct, while control cells received a non-targeting siRNA. **A:** Control and *Paqr5*-silenced MBECs were seeded in equal numbers (75,000 cells/cm^2^) on E-plates having golden electrodes in the bottom of the wells. Impedance of the cell layers represented as the cell index was measured every 15 minutes. Graph shows one representative of N = 2 independent experiments, each performed in sextuplicates. **P ≤ 0.01 compared to control cells receiving the non-targeting siRNA (areas under curves compared with Student’s *t*-test). B: Control and *Paqr5*-silenced MBECs were seeded in equal numbers (75,000 cells/cm^2^) on semipermeable filter inserts. After reaching confluence, the monolayers were supplied with hydrocortisone (HC) and cAMP (CPT-cAMP + Ro 20-1724) to induce maturation of TJs. TEER was measured every 30 minutes with the cellZscope instrument during both the growth and maturation phases. Graph shows one representative of N = 2 independent experiments, each performed in triplicates. ***P ≤ 0.001 compared to control cells receiving the non-targeting siRNA (areas under curves compared with Student’s *t*-test). C: Permeability of sodium fluorescein through control and *Paqr5*-silenced MBEC monolayers was performed for 30 minutes, 24 hours after addition of HC and cAMP. Graph shows one representative of N = 2 independent experiments, each performed in triplicates. * P ≤ 0.05 compared to control cells receiving the non-targeting siRNA (Student’s *t*-test). D: Total protein was extracted for western blot from control and *Paqr5*-silenced cells. Left panels show representative blots, β-actin was used as loading control. Graph represents average ± SD from N = 3 independent experiments. ***P ≤ 0.001 compared to control cells receiving the non-targeting siRNA (Student’s *t*-test). E: Claudin-5 immunofluorescence staining was performed on confluent, control and *Paqr5*-silenced MBEC layers. One representative confocal micrograph (maximum intensity projection of *z*-stacks) of N = 2 independent experiments is shown. Arrows indicate disruption in the continuity of the staining.

To more precisely assess the permeability of TJs to ions and small molecules, TEER and sodium fluorescein penetration were measured. In *Paqr5*-silenced cells, a slower increase in TEER was observed with a lower plateau suggesting impaired TJ maturation (**Fig. 6 B**). Accordingly, the sodium fluorescein permeability of monolayers formed by *Paqr5*-silenced cells was significantly higher than that of the control (**Fig. 6 C**). As a potential molecular mechanism underlying TJ opening, moderately yet significantly decreased claudin-5 levels were observed in *Paqr5*-downregulated cells (**Fig. 6 D**), which was also reflected by fainter and partially disrupted claudin-5 junctional immunostaining (**Fig. 6 E**).

Impaired barrier integrity may enhance the migration of tumour cells from the apical (luminal) side to the basolateral (brain) side of the endothelium. To test this hypothesis, we performed *in vitro* transendothelial migration assays using control and *Paqr5*-silenced MBECs cultured on large pore-size filter inserts and 4T1-tdTomato cells. The assay was conducted for 48 hours to allow sufficient time for cancer cell migration while ensuring that unchallenged endothelial monolayers remained intact. At this time point, the majority of tumour cells were either attached to the apical surface of, or intercalated among, control endothelial cells. On the other hand, clusters of transmigrated tumour cells could be detected on the basolateral side of the *Paqr5*-silenced endothelial monolayers, as shown in the orthogonal views of the confocal micrographs (**Fig. 7 A**). After removing endothelial and tumour cells from the upper sides of the filters, we quantified the cancer cells that had migrated through the endothelium and reached the bottom surface of the porous membrane (Wilhelm et al., 2011). In comparison to control MBECs, we observed significantly more cancer cells migrated through *Paqr5*-silenced endothelial monolayers confirming enhanced transmigration in the absence of the receptor (**Fig. 7 B**).

**Figure 7.**
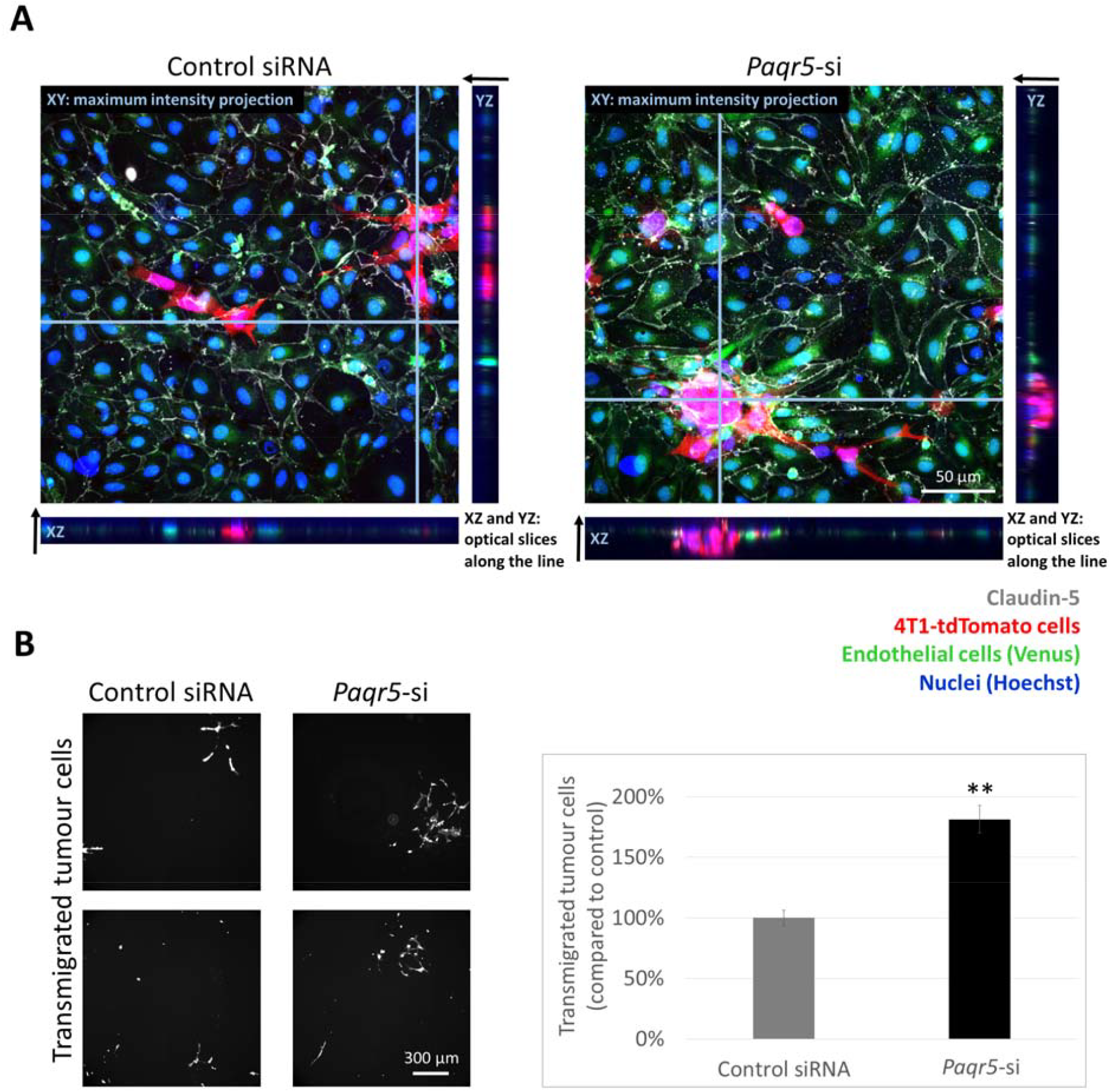
Enhanced transmigration of TNBC cells across brain endothelial monolayers with reduced *Paqr5* expression. Control and *Paqr5*-silenced MBECs were cultured on large pore-size filter inserts until confluence. 4T1-tdTomato cells were placed in the top compartment at a density of 10^5^ cells/cm^2^ and left for 48 hours. A: Claudin-5 immunofluorescence was performed. One representative confocal micrograph is shown for both control and *Paqr5*-silenced MBEC monolayers 48 hours after addition of the tumour cells. XY images show maximum intensity projection of *z*-stacks, while XZ and YZ views represent single optical slices along the light blue lines on the XY images. Arrows indicate bottom-to-top directions in the orthogonal sections. B: After removal of the cells from the top side of the filters, tumour cells that had migrated through the endothelial monolayer and the pores of the filter were counted. In the left panels, two representative epifluorescence images are shown for both conditions. Graph on the right represents average ± SD from N = 2 independent experiments, each performed in triplicates. Please note that the difference may be underestimated, as clusters of cells migrating through *Paqr5*-silenced MBEC monolayers could not always be optically resolved into individual cells. **P ≤ 0.01 compared to control cells receiving the non-targeting siRNA (Student’s *t*-test).

Moreover, we observed that *Paqr5*/mPRγ might also modulate other endothelial functions including cell migration and angiogenesis. This was indicated by the much slower repair of the scratched monolayer in *Paqr5*-silenced cells in the wound closure assay (**Suppl. fig. 5 B, C**), as well as by inhibited tube formation in EV-encountered and *Paqr5*-silenced MBECs (**Suppl. fig. 6**).

Collectively, these results indicate that PAQR5 is essential for maintaining and restoring cerebral endothelial monolayer and barrier integrity, as well as for preventing cancer cell migration from the luminal to the brain side.

## Discussion

EVs regulate multiple stages of tumourigenesis, including organ-specific metastasis, through shaping the pre-metastatic microenvironment and reprogramming recipient cells to facilitate cancer progression (Kalluri and McAndrews, 2023; Urabe et al., 2021). During the development of cerebral secondary tumours, EVs can cross the BBB (Morad et al., 2019), influencing immune responses and metabolism, as well as remodelling the vasculature (Li et al., 2024). However, little is known about how EVs contribute to the migration of cancer cells across brain vessel walls that form the BBB, which is a key step in the development of brain metastases (Wilhelm et al., 2018). As described in mouse models, tumour cells spend several days arrested in cerebral microvessels before being able to complete diapedesis into the brain (Kienast et al., 2010; Lorger and Felding-Habermann, 2010). The prolonged retention of metastatic cells within capillary lumens is characteristic only of the CNS (Paku et al., 2000). During these 2–5 or more days, arrested cells induce several changes in the endothelium, including vessel obstruction and blebbing (Haskó et al., 2019), as a result of communication between the two cell types. One of the channels through which arrested or distant tumour cells may influence the brain endothelium is the secretion of EVs, which are subsequently internalized by host cells. Metastatic breast cancer-released EVs may open the TJs – the main components of barrier properties of brain endothelial cells – through miR-105- or miR-181c-dependent mechanisms (Tominaga et al., 2015; Zhou et al., 2014).

Here we show that EVs from TNBC cells open the BBB via miR-146a-5p- and TGF-β1-mediated downregulation of endothelial *Paqr5*/PAQR5 leading to disruption of the TJs, particularly repression of claudin-5. Our results were obtained primarily using a highly relevant *in vitro* mouse model – primary brain endothelial cells – which retain neural environment-induced barrier properties, as reflected by continuous TJs at cell borders and high TEER values. In addition, we partially confirmed our findings in human cell cultures and in a mouse brain metastasis model.

Using a screening assay followed by qPCR validation, we first identified elevation of miR-146a-5p levels in brain endothelial cells encountering EVs released by TNBC cells. The miR-146 family comprises two closely related members, miR-146a and miR-146b, which differ by only two nucleotides. Both generate the mature −5p strand as the bioactive guide strand and are highly conserved across species (Paterson and Kriegel, 2017).

MiR-146a is a key miRNA that functions as a negative feedback regulator of NF-κB, thereby playing an important role in inflammatory and malignant processes (Zhao et al., 2011). MiR-146a-5p is among the most abundant plasma miRNAs, which is upregulated in sepsis and functions to downregulate the IRAK1 protein (Wang et al., 2021). It is also overexpressed in the serum and tissue samples of non-small cell lung cancer patients and plays a pro-oncogenic role via suppression of TRAF6 (Liu et al., 2020). MiR-146a has been shown to mitigate myocardial ischaemia–reperfusion injury through decreasing MED1/TRAP220 expression (Zhang et al., 2019) and it plays a pivotal role in a range of neurodegenerative and neuro-inflammatory disorders as well (Fan et al., 2020; Rawat et al., 2025). In the brain, it is secreted by microglia and actively transferred via EVs to diminish neurogenesis in depression (Fan et al., 2022), while in glioblastoma (GBM), inflammatory M1 macrophages release EVs containing miR-146a-5p that downregulate TRAF6 and IRAK1 in GBM cells, thereby modulating tumour-associated inflammatory signalling (Xu et al., 2023).

In breast cancer, the expression of miR-146a and miR-146b has been linked to the basal-like/TNBC subtype (Garcia et al., 2011), to stem cell-like properties and therapy resistance (Tordonato et al., 2021), as well as to activation of cancer-associated fibroblasts (Yang et al., 2020). Conversely, miR-146a has been shown to decrease the migratory capacity of TNBC cells (Bhaumik et al., 2008; Wang et al., 2013), and its repression has been associated with poor treatment outcomes, recurrence, and metastasis (Panoutsopoulou et al., 2023), underlying the potential double role of miR-146a as both an oncogenic and a tumour suppressor miRNA (Iacona and Lutz, 2019). Among brain metastatic tumours, miR-146a-5p is highly expressed in cerebral metastases of melanoma. EV-mediated transfer of miR-146a-5p from melanoma cells to astrocytes suppresses NUMB within the Notch signalling pathway, leading to the activation of tumour-promoting cytokines (Rigg et al., 2023). However, the involvement of miR-146a-5p and its interplay with the TGF-β1 pathway in the development of breast cancer brain metastases has previously been unknown.

In our experimental setup, the EV origin of miR-146a has not been definitively demonstrated, although it is suggested by its high abundance in EVs. Nevertheless, we cannot exclude the possibility that EVs induce the upregulation of endogenous miR-146a in endothelial cells. As components of a ubiquitous signalling pathway (Saba et al., 2014), endothelial miR-146a and miR-146b accumulate during the later stages of the inflammatory response to limit excessive inflammation (Cheng et al., 2013); however, their expression may also be upregulated under other conditions.

We observed downregulation of five miR-146a target genes in brain endothelial cells upon exposure to EVs released by TNBCs, as well as after transfection with synthetic miR-146a. Four of these genes are well-known miR-146a targets implicated in diverse diseases, encoding the MED1, NUMB, TRAF6, and IRAK1 proteins, as indicated above. As a less well-characterized target, *Paqr5* within the brain is almost exclusively expressed in endothelial cells. Moreover, this target, but not the other four, was also repressed by another EV cargo, TGF-β1, which exerted a more robust effect on this gene than miR-146a. Although we have not studied the interaction of EV-delivered TGF-β1 with its receptor on the endothelial cells, prior studies suggest that latent TGF-β1 on the surface of EVs is activated within the endosomal compartment of recipient cells through acidification, resulting in prolonged signalling via receptors internalized into endocytic membranes (Shelke et al., 2019).

Tumour-secreted TGF-β1 has previously been shown to alter the integrity of brain endothelial TJs through the induction of endothelial–mesenchymal transition (Krizbai et al., 2015); however, the involvement of PAQR5/mPRγ has not been linked to this phenomenon.

Nevertheless, PAQR5 has been implicated in epithelial–mesenchymal transition (Yang et al., 2025), a closely related process in which TGF-β1 is a key coordinator (Wang et al., 2023). Moreover, suppression of PAQR5 expression by TGF-β1 has also been reported previously in another tumour type (Tao et al., 2022). In addition, miR-146a-5p was shown to regulate endothelial permeability by directly targeting Junctional Adhesion Molecule C (JAM-C) (Hu et al., 2020). All these data support our observations that miR-146a, TGF-β1, PAQR5 and the TJs are mechanistically connected.

Nevertheless, regulation of brain endothelial PAQR5 by metastatic cells represents a previously unknown mechanism, which is particularly relevant given that this membrane progesterone receptor is predominantly and selectively expressed in endothelial cells among cells of the CNS. In addition, regulation of the BBB by PAQR5 represents a novel mechanism that may also be essential under physiological conditions. The BBB is a unique function of the cerebral endothelium through which these cells strictly regulate the entry of substances and cells into the brain. By sealing the intercellular cleft, continuous TJs, and especially claudin-5, form the structural basis of the barrier (Hashimoto et al., 2023). TJ protein complexes exhibit high dynamism and undergo modification in response to multiple physiological and pathological stimuli (Zihni et al., 2016). The protective effect of progesterone on the BBB under diverse pathological conditions, mediated partly by restoration of claudin-5 levels, has been well established (Espadín et al., 2025; Ishrat et al., 2010); however, the involvement of nuclear and membrane receptors has not yet been clarified. Here we show that Org, an agonist of mPRα that is also likely to act on the structurally related mPRγ as well, increases claudin-5 expression in brain endothelial cells in a similar manner to progesterone. These seven-transmembrane proteins most likely regulate TJ proteins through modulation of cAMP levels and the MAPK signalling pathway (Mauvais-Jarvis et al., 2022); however, further studies are needed to understand the exact mechanisms.

The BBB is also responsible for restricting metastatic cells from migrating from the vessel lumen into the brain parenchyma (Wilhelm et al., 2013). Metastatic breast cancer cells are able to use both the transcellular and paracellular pathways, being able to migrate through individual endothelial cells and through the TJs (Herman et al., 2019). This novel mechanism of PAQR5/mPRγ downregulation and subsequent reduction of claudin-5 levels in the TJs favours paracellular diapedesis of the tumour cells.

Interestingly, the migratory and angiogenic properties of cerebral endothelial cells were also inhibited upon reduction of PAQR5 expression. Although it may initially seem counterintuitive that tumour cells would inhibit angiogenesis, it should be noted that vascularization of breast cancer brain metastases primarily relies on vascular co-option rather than angiogenesis (Kienast et al., 2010). More importantly, our analyses focused on early stages of brain metastasis formation, specifically prior to and during transmigration across the BBB, when tumour cell-derived signals, including EVs, reach the brain endothelium from the luminal side. At this stage, with individual tumour cells localized within the vessel lumen, endothelial responses to these signals may play a critical role in determining whether tumour cells successfully penetrate the BBB. A limitation of our study is that this incipient step of brain metastasis formation is practically impossible to investigate in human patients, as current clinical imaging techniques lack the resolution to detect single tumour cells. However, our use of human cells, together with the evolutionary conservation of miR-146a across species, supports the translational relevance of our findings. Further studies are nevertheless required to elucidate the effects of abluminally localised tumour cells on PAQR5 expression in endothelial cells of the blood– tumour barrier in established metastatic lesions.

In conclusion, we have identified a novel mechanism through which TNBC cells impair the BBB to facilitate their migration into the brain. Tumour cell-derived EVs induce miR-146a-5p- and TGF-β1-mediated downregulation of PAQR5 in cerebral endothelial cells, leading to repression of claudin-5 in the TJs. Consequently, TJ opening enhances the transendothelial migration of cancer cells (**Suppl. fig. 7**). Additional studies are warranted to assess the therapeutic potential of PAQR5 modulation for BBB protection in metastatic and other CNS disorders.

## Statements

### Data availability statement

The data supporting the findings of this study have been deposited in the ARP repository (https://repo.researchdata.hu/) and will be made freely available upon acceptance of the manuscript for publication.

### Ethics approval statement

All animals were housed and handled in accordance with established ethical standards, and all experimental procedures were approved by the institutional animal care committee and the Regional Animal Health and Food Control Station of Csongrád-Csanád County (permit numbers: XVI./764/2018. and XVI./2161/2024.).

## Acknowledgments

We acknowledge the contributions of Kinga Mészáros-Molnár and Ádám Mészáros to the preliminary experiments of the present study. We also acknowledge András Kincses for his work on the dynamic light scattering measurements. Microscopy imaging has been partly performed in the Cellular Imaging Laboratory of the HUN-REN Biological Research Centre with the help of Gábor Steinbach. We gratefully acknowledge the receipt of FVB/Ant:TgCAG-YFP mice from Ferenc Erdélyi (Institute of Experimental Medicine, Budapest, Hungary).

## Authors’ contributions

I.A.K. and I.W. designed the research study; C.F., T.D., D.V., and A.L. performed research; C.F., A.G.V., M.K., A.E.F., I.A.K. and I.W. analysed the data; I.W. and I.A.K. supervised research; I.W. drafted the manuscript; all authors approved final version.

