## Supplemental file for "PAQR5 Membrane Progesterone Receptor Regulates the Blood–Brain Barrier during Brain Metastasis Formation"

### SUPPLEMENTARY MATERIAL

#### Supplementary methods

##### Characterization of EV size and morphology

The size distribution of the samples was determined by dynamic light scattering using a Zetasizer Nano ZS instrument (Malvern Panalytical, Malvern, UK). All measurements were carried out in PBS at room temperature, with three measurements performed for each sample using automatic resolution and instrument parameters calculated by the driver software.

Atomic force microscopy imaging was carried out with an MFP-3D instrument (Asylum Research, Santa Barbara, CA USA) mounted on an Axiovert 200 optical microscope (Zeiss) using the Igor Pro 6.32A driving software (WaveMetrics, Portland, OR USA), as described previously (Fazakas et al., 2021). BL-RC150VB-HW gold-coated silicon nitride cantilevers were used, featuring a V-shaped tip with a 30-nm radius of curvature, a nominal spring constant of 50 pN/nm, and a resonant frequency of 37 kHz (Olympus, Tokyo, Japan). Images were recorded in PBS after immobilization of the EVs onto a glass surface using glutaraldehyde.

##### Supplementary cell culture methods

Primary mouse astrocytes were isolated from two-day-old pups by mechanical dissociation of the brain tissue. Cells were cultured in poly-L-lysine-coated dishes in DMEM supplemented with 10% FBS (Thermo Fisher Scientific) and used after three weeks.

D3 cells (hCMEC/D3; received from Prof. P.-O. Couraud at Inserm, Paris, France) were maintained in rat-tail collagen-coated dishes in EBM-2 medium supplemented with EGM-2MV bullet kit (Lonza).

Transfection of D3 cells with the synthetic miR-146a-MIMIC was performed similarly to the procedure applied for 4T1-tdTomato cells. TGF-β1 and SB-431542 were added to the cells in serum-free EBM-2 medium. Org (Org OD 02-0 or 10-ethenyl-19-norprogesterone; MedChemExpress, Princeton, NJ USA) and Prog (progesterone) were applied onto cell cultures at a concentration of 100 nM in serum-free EBM-2 medium.

The brain-seeking BrM2 human TNBC cells (MDA-MB-231-BrM2; obtained from Prof. J. Massagué at Memorial Sloan Kettering Cancer Center, New York, NY USA) were cultured in DMEM supplemented with 10% FBS (Thermo Fisher Scientific). The cells were authenticated by STR analysis (Microsynth).

All cultures were regularly verified for the absence of mycoplasma contamination using the MycoAlert kit (Lonza).

Collection and characterization of EVs released by BrM2 were performed similarly to those of 4T1-tdTomato-derived EVs, as described in detail in (Fazakas et al., 2021).

#### Wound healing assay

Confluent MBEC monolayers were scratched with a pipette tip, washed twice with serum-free media, and incubated in their normal culture media for 48 hours to close the wound. Phase-contrast images of the cells were taken with an Eclipse TE2000 U inverted microscope at the beginning and the end of the experiment. The wound surface was measured with the Image J image processing program.

#### Tube formation assay

MBECs were seeded at a density of 75,000 cells/cm$^2$ in serum-free OptiMEM medium (Thermo Fisher Scientific) into chambered coverslips for 3D cultures (Ibidi, Gräfelfing, Germany) preloaded with 10 µl/well growth factor reduced Matrigel (Corning). After 7 hours, phase-contrast images of the cells were taken with an Eclipse TE2000 U inverted microscope. Images were analysed with the Angiogenesis Analyzer plugin of the Image J software (Carpentier et al., 2020).

### Supplementary tables

#### Supplementary table 1. Synthetic RNA sequences used.

| **Name** | **RNA sequence** | **Source** |
|---|---|---|
| miR-146a-MIMIC (hsa-miR-146a-5p miRCURY LNA miRNA Mimic) | UGAGAACUGAAUUCCAUGGGUU | Qiagen, GeneGlobe ID: YM00472124 |
| Paqr5-si (Paqr5_NM_028748.2_stealth_319) | CAGCCUCUUCCAAAUGACCAAUGAG | Thermo Fisher Scientific |
| Control siRNA (NM_028748.2_stealth_control_319) | CAGUUCUAACCAGUAAACCUCCGAG | Thermo Fisher Scientific |

#### Supplementary table 2. Predesigned miRCURY LNA miRNA PCR assays.

| **Name** | **Source (Qiagen, cat. no. 339306)** |
|---|---|
| miR-146a-5p (hsa) | GeneGlobe ID: YP00204688 |
| SNORD65 (mmu) | GeneGlobe ID: YP00203910 |
| SNORD48 (hsa) | GeneGlobe ID: YP00203903 |
| UniSp6 | GeneGlobe ID: YP00203954 |

#### Supplementary table 3. Primers used for detecting mRNA expression.

| **Name** | **Forward primer (5' → 3')** | **Reverse primer (5' → 3')** |
|---|---|---|
| Paqr5 (m) | AAGCTACCTTGATGTCCCTGGT | CTCCACACAAAGAACCAGAAGGG |
| Med1 (m) | CTCCTCCACTCGTGAAAGGT | ACGACCCTCTTCTCCATTACT |
| Traf6 (m) | CAGCGTGTACGATCGGGTT | AAGACCCAAGTTTCCGTGCC |
| Irak1 (m) | CTCAGATCCCAGGAACAGGTAT | GAACCTGGAGTCAAGTGCCA |
| Numb (m) | GGATGGGCTCAGAGTTGTGG | TCAGTCTTTCCCCCGTGTCT |
| Paqr6 (m) | AGCACAGGCATCTCTCCTAC | ACAGGAAGTACCAGGTGGG |
| Paqr7 (m) | CTCAGCCCATCAGAAGTTTTCC | GCTGAAACAGTGTGCGGAAG |
| Paqr8 (m) | GGTCGGGGAGGCTCAGAAAG | GACCTGAGGAACTCGAACACC |
| Rplp0 (m) | AGATTCGGGATATGCTGTTGGC | TCGGGTCCTAGACCAGTGTTC |
| PAQR5_a (hu) | TGCTGCCCTTCTGGTTCTTT | GTTGACGGCACCATAGTCCA |
| PAQR5_b (hu) | GGACTCCCTCCCCATCTTCT | CTGGTGACTGTGACCGATGT |

| | | |
|---|---|---|
| Tgfb1 | CACCGGAGTTGTGCGGCAGT | TGCCGCACGCAGCAGTTCTT |
| GAPDH | GTGAAGGTCGGTGTCAACG | GTGAAGACGCCAGTAGACTC |

m = mouse, hu = human

#### Supplementary table 4. List of primary antibodies.

| Method | Primary antibody | Host | Dilution | Source | Cat. no. |
|---|---|---|---|---|---|
| IF | anti-Paqr5 | rabbit | 1:100 | Thermo Fisher Scientific | PA5-106846 |
| IF | anti-ZO1 | rabbit | 1:100 | Thermo Fisher Scientific | 61-7300 |
| IF | anti-CD31 | goat | 1:200 | Bio-Techne | AF3628 |
| IF | anti-RFP | rabbit | 1:100 | Rockland Immunochemicals (Limerick, PA USA) | 600-401-379RTU |
| IF, WB | anti-claudin-5 | rabbit | 1:300 | Thermo Fisher Scientific | 34-1600 |
| WB | anti-β-actin | mouse | 1:10000 | Merck/MilliporeSigma | A5441 |
| WB | anti-CD81 | mouse | 1:300 | Bio-Techne | NB100-65805 |
| WB | anti-Hsp70 | mouse | 1:1000 | BD Transduction Laboratories (Franklin Lakes, NJ USA) | 610608 |
| WB | anti-Alix | rabbit | 1:300 | Bio-Techne | NB100-65678 |
| WB | anti-GRP94 | mouse | 1:1000 | Cell Signaling (Danvers, MA USA) | 20292 |

IF = immunofluorescence, WB = western blot

#### Supplementary table 5. List of secondary antibodies.

| Method | Secondary antibody | Host | Dilution | Source | Cat. no. |
|---|---|---|---|---|---|
| IF | anti-rabbit IgG, Alexa Fluor 647 | donkey | 1:500 | Thermo Fisher Scientific | A32795 |
| IF | anti-rabbit IgG, Alexa Fluor Plus 594 | donkey | 1:500 | Thermo Fisher Scientific | A32754 |
| IF | anti-goat IgG, Alexa Fluor 488 | donkey | 1:500 | Thermo Fisher Scientific | A32814 |
| WB | anti-rabbit IgG, HRP | goat | 1:3000 | Cell Signaling | 7074 |
| WB | anti-mouse IgG, HRP | goat | 1:3000 | Thermo Fisher Scientific | G21040 |

HRP = horseradish peroxidase

### Supplementary figures

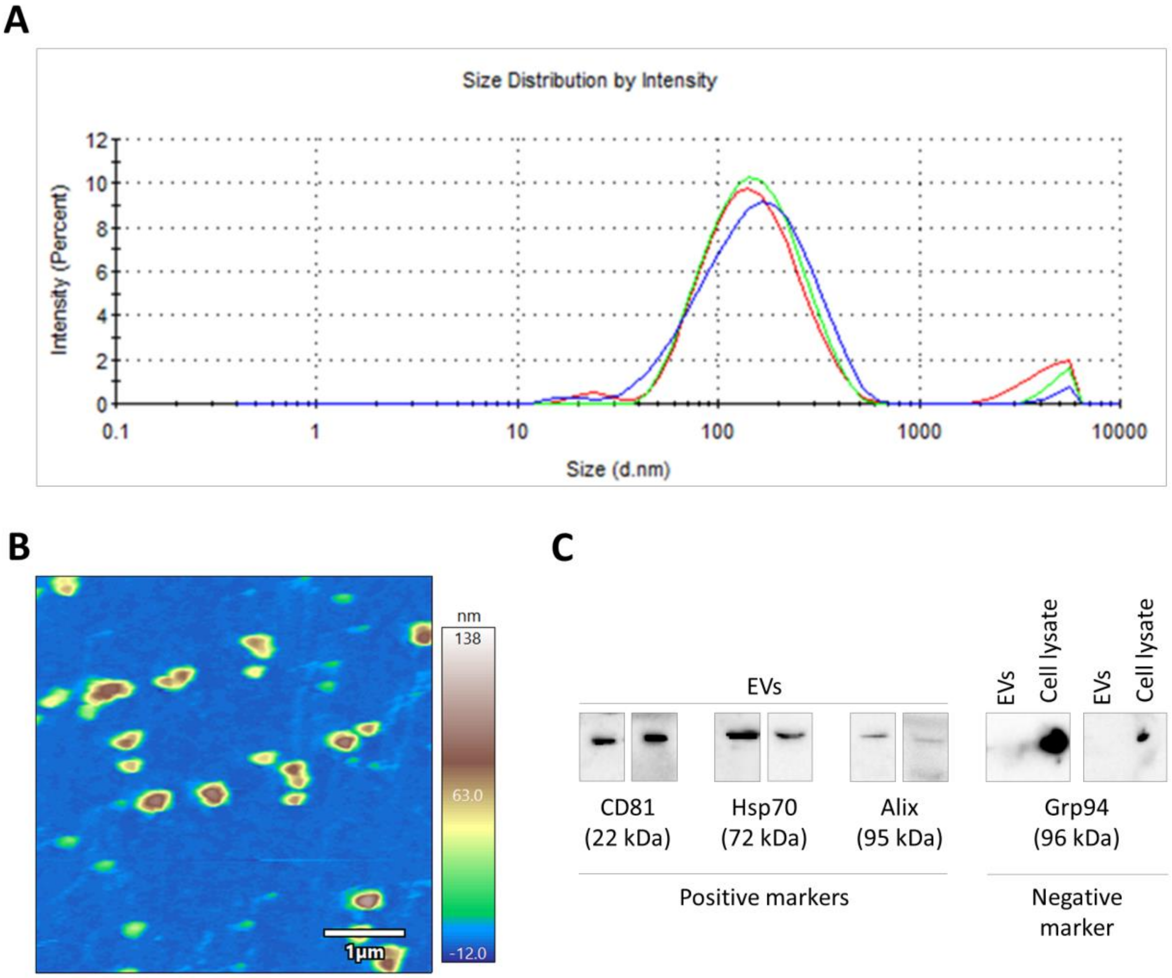


**Supplementary figure 1. Characterization of EVs isolated from mouse TNBC cells.** EVs were isolated from the supernatants of 4T1-tdTomato cells cultured for 24 hours in media containing exosome-depleted FBS. EVs were collected into PBS and stored at –20°C. **A:** Size distribution of the EVs was determined using dynamic light scattering. Representative data showing three consecutive measurements of the same sample are shown in the image. Average diameter is 139.4 nm. **B:** Image shows a representative atomic force microscopy height image of the EVs immobilized onto a glass surface. **C:** Expression of CD81, Hsp70, Alix, and Grp94 proteins was assessed by western blot using 20 µg total protein per lane extracted from the EVs. Each panel shows representative blots from two independent samples. For the negative marker Grp94, 4T1-tdTomato cell lysates were used as positive controls.

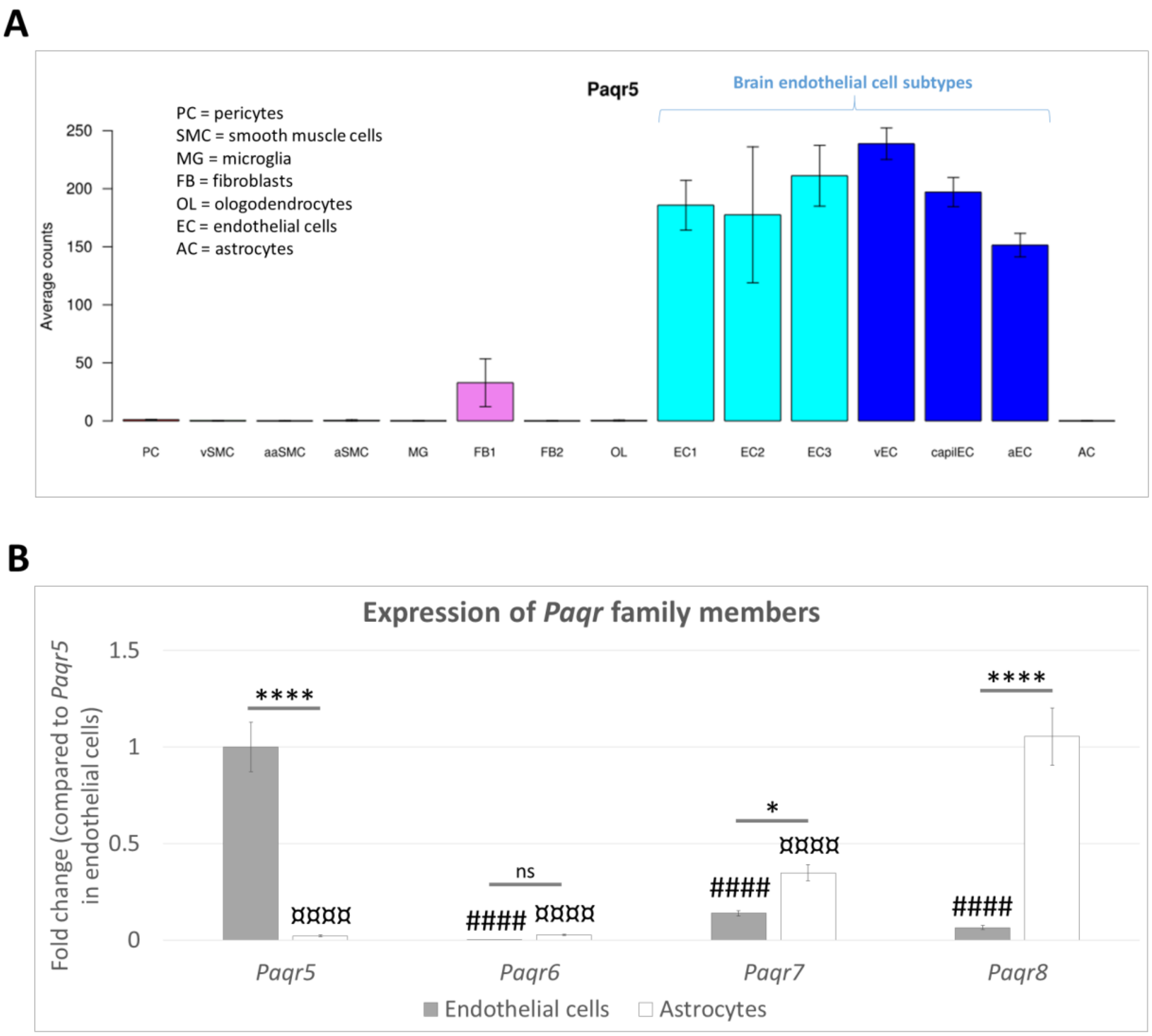


**Supplementary figure 2. Expression of *Paqr* family members in cells of the neurovascular unit. A:** Data on *Paqr5* expression in mouse brain vascular and glial cells were extracted from https://betsholtzlab.org/VascularSingleCells/database.html. **B:** Expression of *Paqr* family members in MBECs and primary mouse astrocytes was assessed by qPCR. Data were normalized to *Rplp0* expression and compared to endothelial *Paqr5* levels. Graph represents average ± SD, n = 3. $^{*}P \le 0.05$, $^{****}P \le 0.0001$, ns = not significant (astrocytes compared to endothelial cells), $^{\#\#\#\#}P \le 0.0001$ (compared to endothelial *Paqr5* expression), ¤¤¤¤P ≤ 0.0001 (compared to astrocytic *Paqr8* expression) (ANOVA and Tukey *post-hoc* test).

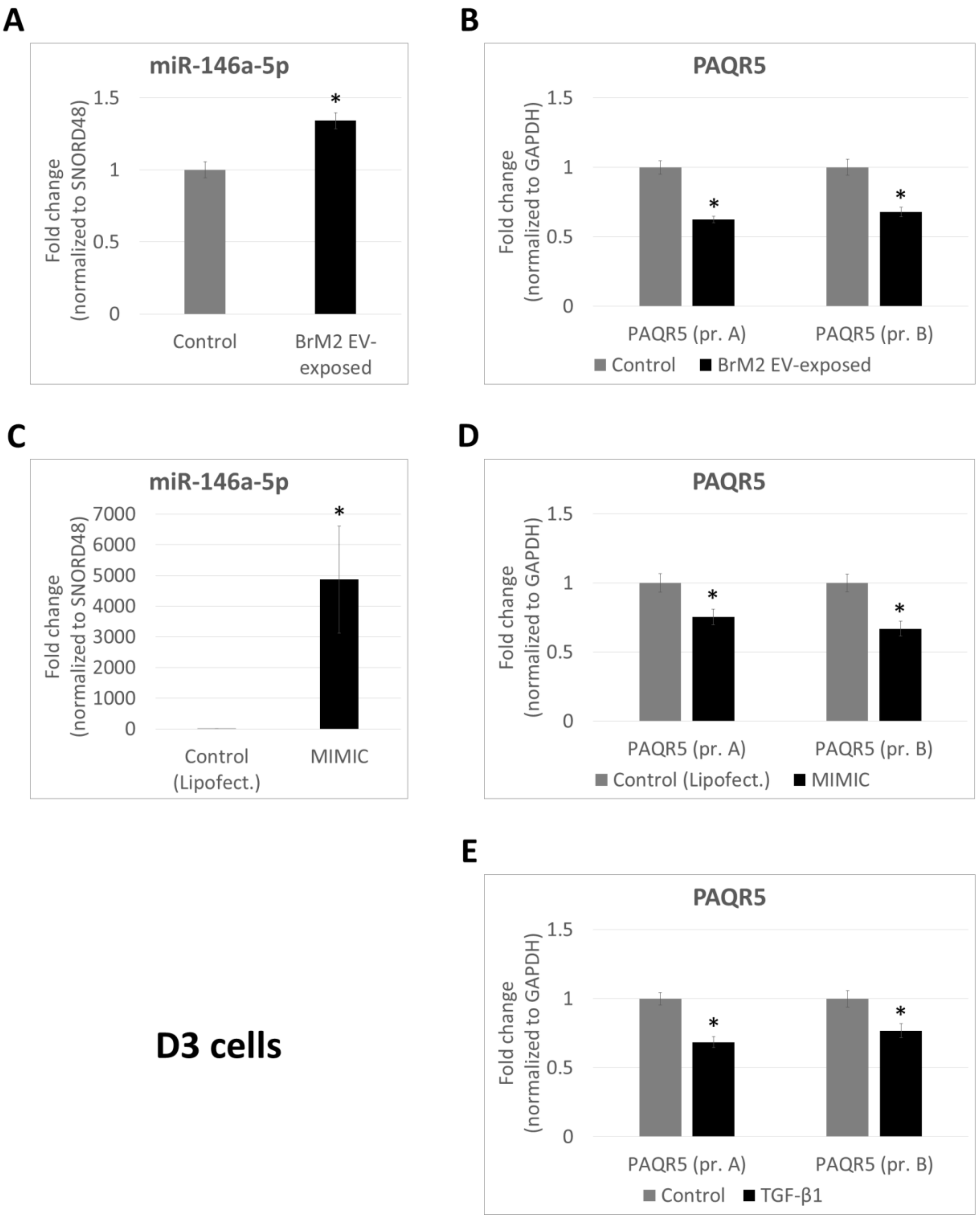


**Supplementary figure 3. Regulation of PAQR5 by miR-146a-5p and TGF-β1 in human brain endothelial cells. A, B:** D3 cells were exposed to BrM2-derived EVs in a dose of 35 µg EV-protein/ml. After 24 hours, total RNA was isolated and miR-146a-5p (**A**) or PAQR5 (**B**) levels were determined by qPCR. For the PAQR5 gene, two different primers (pr. A and pr. B) were used. Graph represents average ± SD from N = 2–4 independent experiments, each performed in triplicates. $^{*}P \leq 0.05$ compared to control (Student's *t*-test for each primer individually). **C, D:** D3 cells were transfected twice with 5 nM miR-146-MIMIC using Lipofectamine, while control cells received the transfection agent only. 24 hours after removal of the transfection reagents, total RNA was isolated and miR-146a-5p (**C**) or PAQR5 mRNA (**D**) levels were determined by qPCR. Graph represents average ± SD from N = 2 independent experiments, each performed in triplicates. Graph represents average ± SD from N = 2 independent experiments, each performed in triplicates. $^{*}$P ≤ 0.05 compared to control (Student's *t*-test for each primer individually). **E:** D3 cells were exposed to 10 ng/ml TGF-β1. After 24 hours, total RNA was isolated and PAQR5 mRNA levels were determined by qPCR. Graph represents average ± SD from N = 2 independent experiments, each performed in triplicates. $^{*}$P ≤ 0.05 compared to control (Student's *t*-test for each primer individually).

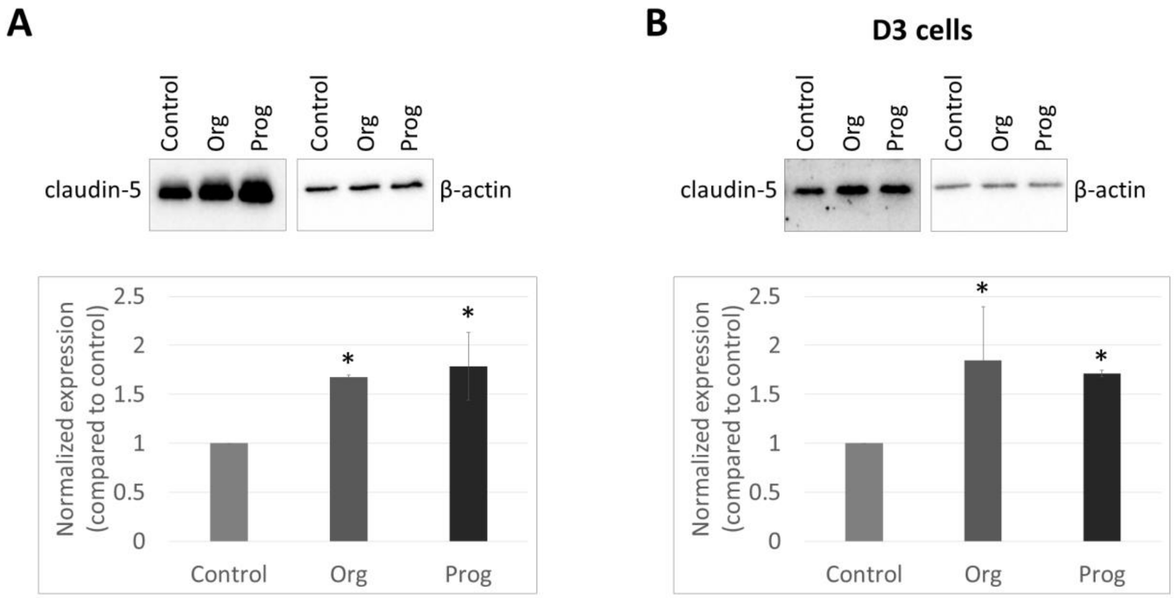


**Supplementary figure 4. Enhanced barrier properties in response to progesterone signalling in mouse and human brain endothelial cells. A:** Total protein was extracted for western blot from MBECs treated for 24 hours with the mPR agonist Org OD 02-0 (Org; 100 nM) or progesterone (Prog; 100 nM). Top panels show representative blots, β-actin was used as loading control. Graph represents average ± SD from N = 2 independent experiments. $^{*}$P ≤ 0.05 compared to control cells (ANOVA and Tukey *post-hoc* test). **B:** Total protein was extracted for western blot from D3 cells treated for 24 hours with Org or Prog. Top panels show representative blots, β-actin was used as loading control. Graph represents average ± SD from N = 3 independent experiments. $^{*}$P ≤ 0.05 compared to control cells (ANOVA and Tukey *post-hoc* test).

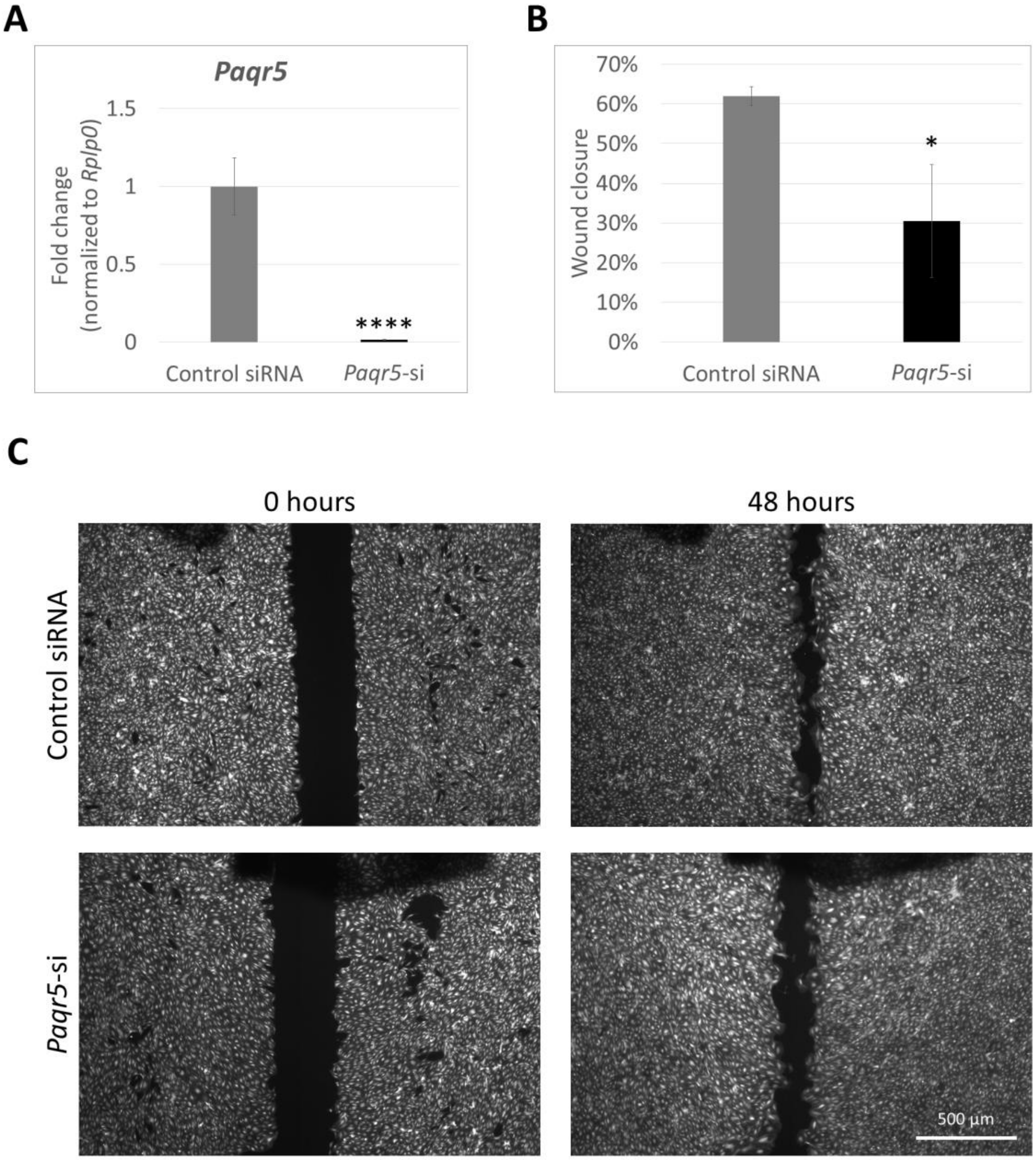


**Supplementary figure 5. Compromised migration properties of *Paqr5*-silenced brain endothelial cells. A:** The efficiency of silencing was verified using qPCR. Graph shows average ± SD from N = 4 independent experiments, each performed in triplicates. $^{****}$P ≤ 0.0001 compared to control cells receiving the non-targeting siRNA (Student's *t*-test). **B, C:** Control and *Paqr5*-silenced MBEC monolayers were scratched with a pipette tip and left for 48 hours to close the wound. Graph in **B** represents average ± SD from N = 3 independent experiments. $^{*}$P ≤ 0.05 compared to control (Student's *t*-test). Representative phase contrast images are shown in **C**.

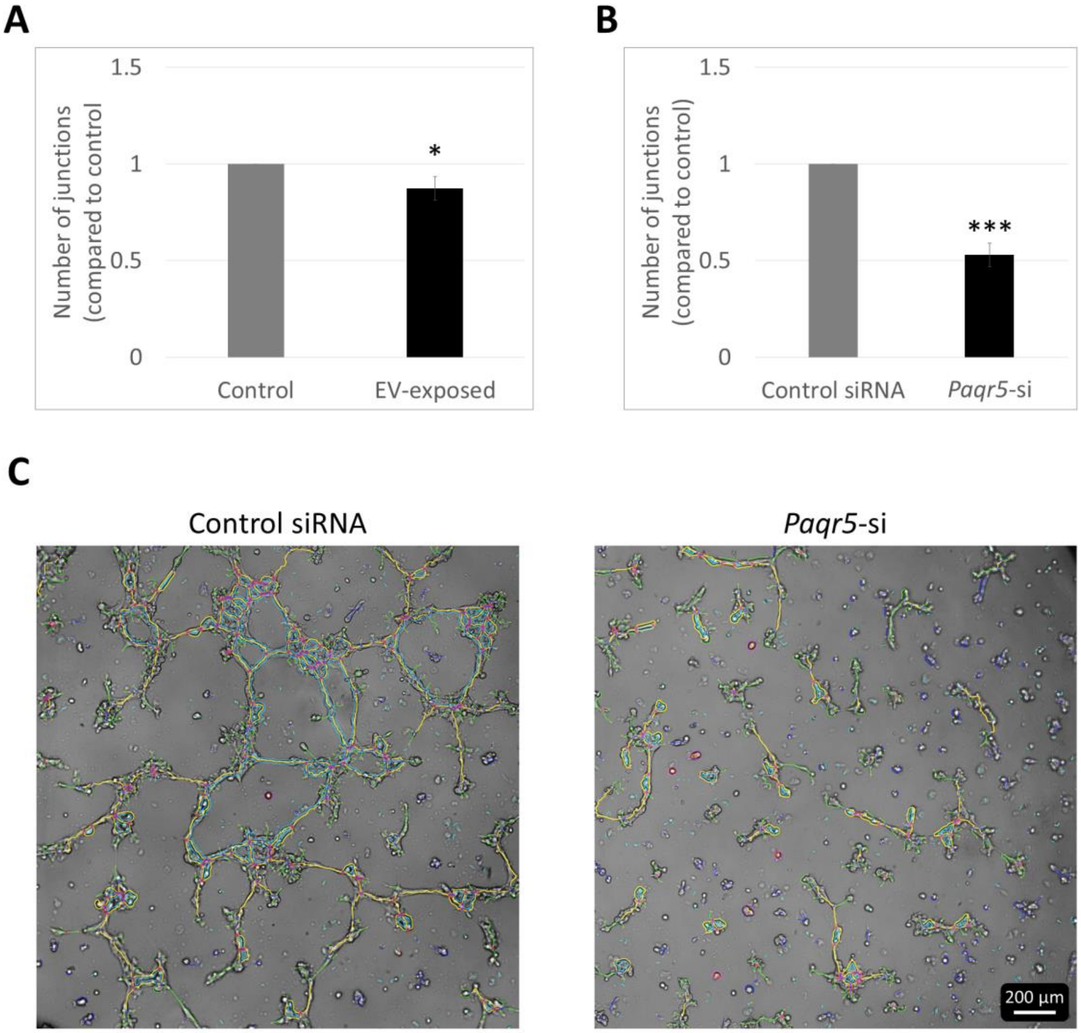


**Supplementary figure 6. Compromised angiogenic properties of *Paqr5*-silenced brain endothelial cells.** MBECs were exposed to 4T1-tdTomato-derived EVs in a dose of 35 µg EV-protein/ml (**A**). Silencing of *Paqr5* in MBECs was performed with a stealth RNA construct, while control cells received a non-targeting siRNA (**B**). Cells were collected and seeded in equal numbers on Matrigel. Phase contrast images of the forming tubes were taken 7 hours later and analysed with Image J. Graphs in **A** and **B** represent average ± SD from N = 3 independent experiments, each performed in triplicates. $^{*}P \leq 0.05$, $^{***}P \leq 0.001$ compared to control (Student's *t*-test). Representative phase contrast images are shown in **C**.

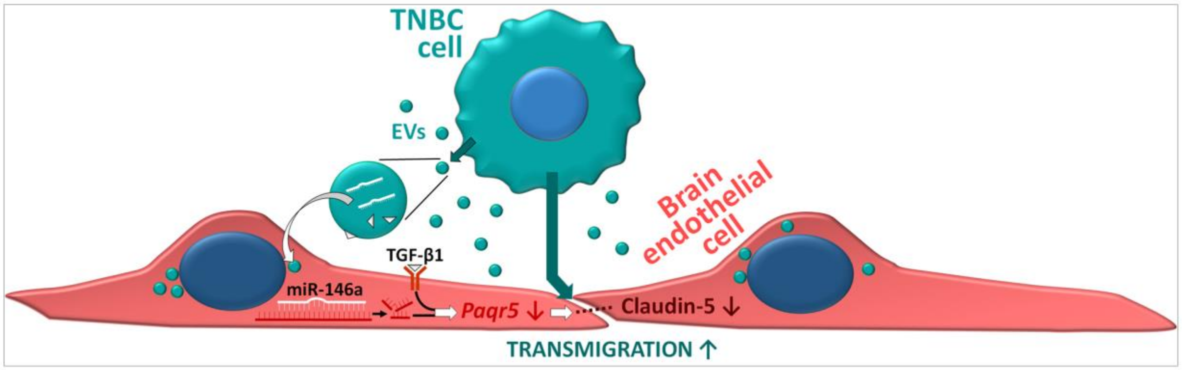


**Supplementary figure 7. Graphical outline of the proposed mechanism.** EVs from TNBC cells induce miR-146a-5p- and TGF-β1-mediated downregulation of *Paqr5*/mPRγ, a membrane progesterone receptor, in BBB-forming endothelial cells. This results in disruption of interendothelial TJs, particularly via repression of claudin-5 protein, thereby promoting enhanced transmigration of cancer cells across the brain endothelium.
